# Conserved core and dynamic periphery NRC helper NLRs underpin immune receptor network evolution across Solanaceae

**DOI:** 10.64898/2026.08.10.743904

**Authors:** Liang-Yu Hou, May Htet Aung, Ian Bien Oloc-oloc, Jiorgos Kourelis, Chih-Hang Wu

## Abstract

Plant nucleotide-binding leucine-rich repeat (NLR) proteins function as intracellular immune receptors that detect pathogen-derived signals and activate defense responses. The NRC (NLR required for cell death) receptor network plays central roles in immunity of solanaceous crops, yet its evolutionary diversification across Solanaceae remains poorly understood. Here, we combined comparative phylogenomics and comprehensive functional complementation assays to investigate the evolution and functional diversification of NRC helper NLRs across nine representative species from diverse genera within the Solanaceae. Phylogenetic analyses resolved 11 NRC helper subfamilies with distinct evolutionary trajectories, revealing a conserved core and dynamic periphery within the NLR receptor network. NRC2, NRC3, and NRC4 were broadly conserved across all examined species, whereas other NRC lineages exhibited degrees of presence–absence polymorphisms, lineage-specific expansion, and rapid diversification. Comparative genomic analyses revealed highly dynamic helper–sensor NLR cluster organization, indicating substantial genomic restructuring during Solanaceae evolution. Functional assays further showed that some NRC subfamilies retained broad compatibility with multiple sensor NLRs despite extensive sequence and genomic divergence, whereas other helpers displayed lineage-specific gains and losses of compatibility, revealing extensive rewiring of helper–sensor functional connections. Together, our study provides a cross-Solanaceae evolutionary and functional atlas of the NRC immune receptor network and demonstrates how a conserved core and dynamic periphery of NRC helper NLRs underpin the evolution of immune signaling specificity across Solanaceae.

## Introduction

Plants rely on intracellular immune receptors to detect pathogen invasion and initiate defense responses. Among these receptors, nucleotide-binding leucine-rich repeat proteins (NLRs) constitute a highly diverse and rapidly evolving family that recognizes pathogen-derived effectors and frequently triggers a localized hypersensitive cell death response (Dodds and Rathjen 2010; Jones et al. 2016; Ngou et al. 2022). NLRs share a conserved tripartite architecture, comprising an N-terminal signaling domain, a central nucleotide-binding domain (NBD), and a C-terminal leucine-rich repeat (LRR) domain (Duxbury et al. 2021). Based on variation in their N-terminal domains, plant NLRs are classified into three major groups: Toll/interleukin-1 receptor (TIR) domain–containing NLRs (TNLs), coiled-coil (CC) domain–containing NLRs (CNLs), and resistance to powdery mildew 8 (RPW8)-like domain–containing NLRs (RNLs) (Duxbury et al. 2021).

Molecular and genetic studies have further classified plant NLRs into three functional modes of action: singletons, pairs, and networks (Adachi et al. 2019; Contreras et al. 2023a). Singleton NLRs can directly or indirectly perceive pathogen effectors and autonomously activate downstream immune responses without requiring additional NLR partners (Contreras et al. 2023a). In contrast, paired NLRs function through a division of labor, in which a sensor NLR specialized for effector recognition cooperates with a helper NLR that mediates immune signaling. Such paired NLRs often physically interact and are frequently encoded as linked gene pairs or clusters within the genome, suggesting coordinated regulation and a history of exclusive co-evolution (Xi et al. 2022). Beyond these binary systems, an increasing number of NLRs operate as components of genetically and functionally interconnected networks, in which multiple sensor and helper NLRs collectively coordinate immune activation. These networked architectures provide both robustness and flexibility, enabling plants to integrate diverse pathogen recognition events into effective defense responses (Wu et al. 2017, 2018; Castel et al. 2019; Saile et al. 2020).

The NLR required for cell death (NRC) family represents a well-characterized example of a networked immune system (Wu et al. 2017, 2018). NRC proteins function as helper NLRs that act downstream of multiple sensor NLRs, including receptors that confer resistance to viruses, bacteria, oomycetes, and nematodes (Wu et al. 2017). Canonical helpers such as NRC2, NRC3, and NRC4 exhibit partial redundancy and collectively mediate immune signaling for a broad repertoire of NRC-dependent sensors of solanaceous plants. For instance, in *Nicotiana benthamiana*, the sensor NLRs Rpi-blb2 and R1 transmit immune signals specifically through NRC4 to trigger cell death (Wu et al. 2017). In contrast, sensor NLRs such as Prf and Rpi-amr1 can signal through either NRC2 or NRC3 (Wu et al. 2016, 2017; Witek et al. 2021; Wu and Kamoun 2021), whereas Rx, Rpi-amr3, Bs2, and Sw5b activate immune responses redundantly via NRC2, NRC3, or NRC4 in *N. benthamiana* (Wu et al. 2017; Chen et al. 2021; Lin et al. 2022). These observations support a model in which three partially redundant NRC helpers form a robust signaling core capable of supporting diverse sensor NLRs in Solanaceae.

Comparative and evolutionary analyses across angiosperms reveal that NRC networks have undergone complex evolutionary trajectories, evolving into lineage-specifically expanded immune signaling networks from an ancestral conserved helper–sensor NLR gene cluster. Phylogenetically, NRC helpers cluster together with NRC-dependent sensor NLRs to form the NRC superclade, a distinct subgroup within CNLs (Wu et al. 2017; Kourelis et al. 2021; Contreras et al. 2023a). The NRC superclade is distributed throughout the asterids and in some Caryophyllales, but is absent from monocots and rosids, suggesting that the ancestral NRC superclade emerged before the divergence of Caryophyllales and asterids (Wu et al. 2017). Phylogenomic analyses further revealed that NRC networks remain relatively simple in Ericales and campanulids, but became highly expanded in lamiids, including Solanaceae (Goh et al. 2024; Sakai et al. 2024). Comparative studies also identified NRC0 as the only NRC helper broadly conserved across asterid lineages, where it frequently forms genetically linked clusters with sensor NLRs, supporting the hypothesis that modern NRC networks originated from an ancestral helper–sensor NLR gene pair (Goh et al. 2024; Sakai et al. 2024).

Beyond this conserved framework, lineage- and organ-specific specialization has further diversified NRC network function. In *Solanum*, NRC6 and its associated clustered sensor NLRs evolved root-specific expression patterns that enable resistance against nematode infection (Lüdke et al. 2025). In *Nicotiana* spp., analyses of natural and synthetic NRC3 alleles demonstrated that subfunctionalization reshapes the genetic architecture and signaling properties of NRC networks (Huang et al. 2024). Supporting the functional significance of this diversification, the helper NLRs NRC8 and NRC9 were identified in pepper (*Capsicum annuum*) and shown to mediate cell death signaling downstream of CaRpi-blb2, a pepper homolog of the wild potato immune receptor Rpi-blb2 from *Solanum bulbocastanum* (Oh et al. 2023). Together, these findings suggest that diversification of NRC repertoires has contributed to the emergence of species-, lineage-, and tissue-specific immune signaling pathways.

At the mechanistic level, structural, biochemical, and cell biological studies have provided major insights into the activation and functions of NRCs following pathogen effector recognition by sensor NLRs. Prior to activation, NRCs exist as inactive homodimers distributed between the cytoplasm and plasma membrane, where intramolecular interactions maintain an autoinhibited resting state (Selvaraj et al. 2024). Recognition of pathogen effectors by upstream sensor NLRs induces conformational rearrangements that relieve this autoinhibition and promote oligomerization of NRCs into higher-order hexameric resistosome complexes (Ahn et al. 2023; Contreras et al. 2023b; Liu et al. 2024; Madhuprakash et al. 2024). Live-cell imaging and biochemical analyses further revealed that activated NRC resistosomes assemble into punctate structures at the plasma membrane that correlate with calcium influx, organelle perturbation, and progression of hypersensitive cell death (Duggan et al. 2021; Chen et al. 2026). Recent advances integrating artificial intelligence–based structural prediction with comparative analyses further revealed that NRC resistosomes exhibit greater structural diversity than previously appreciated, including atypical higher-order assemblies distinct from the canonical hexameric architecture (Toghani et al. 2026b). Together, these findings establish NRCs as dynamic signaling hubs that transition from a quiescent resting state into membrane-associated resistosome complexes that coordinate immune signaling and cell death execution.

Despite these advances, our understanding of NRC evolution remains incomplete in several key aspects. Most functional and evolutionary studies have focused on *Solanum* and *Nicotiana*, leaving NRC diversity in other solanaceous genera largely unexplored. It remains unclear how many NRC subfamilies exist within Solanaceae, how their genomic organization relates to functional specialization, and whether helper–sensor compatibility is broadly conserved or shaped by lineage-specific divergence. Here, we address these questions by reconstructing the evolutionary history and functional landscape of NRC variants across nine solanaceous species spanning multiple genera. Through integrated phylogenomic analyses, we identify 11 NRC helper subfamilies and reveal distinct evolutionary regimes characterized by a conserved core and a rapidly diversifying periphery. Extensive cross-species functional assays uncover both widespread redundancy and pronounced subfunctionalization among NRC helpers, including an unexpectedly broad role for NRC9 beyond the *Solanum* lineage. Together, our findings provide a unified framework linking NRC evolution, genomic organization, and immune function, offering new insights into how NLR networks diversify while maintaining robust disease resistance in Solanaceae.

## Result

### Phylogenomic analyses resolve conserved and lineage-specific NRC helper subfamilies in Solanaceae

To investigate the evolutionary trajectory of the NRC network within Solanaceae, we retrieved the predicted proteomes from nine representative species, including *Petunia inflata* (petunia), *Nicotiana benthamiana*, *Lycium barbarum* (goji berry), *Datura stramonium* (jimsonweed), *Physalis floridana* (lantern fruit), *Capsicum annuum* (pepper), *Solanum melongena* (eggplant), *Solanum lycopersicum* (tomato), and *Solanum tuberosum* (potato), for comparative phylogenomics (Fig. 1A, Table S1). We applied the established NLRtracker pipeline to identify NLR candidates and extracted their NB-ARC domains for multiple sequence alignment using MAFFT (Fig. 1B) (Kuraku et al. 2013; Katoh et al. 2019; Kourelis et al. 2021). We then performed phylogenetic reconstruction using IQ-TREE (Trifinopoulos et al. 2016), and used the resulting tree to identify the NRC superclade of individual plant species (Fig. 1B). The sequences were then combined into a single file. After sequence alignment and trimming, sequences shorter than 200 amino acids were excluded to improve the reliability of phylogenetic inference. The resulting alignment was then used to generate a phylogenetic tree encompassing NRC superclade members from all nine species (Fig. 1C, Table S2, Dataset S1). Consistent with the previously described organization of the NRC network, the combined phylogeny across diverse solanaceous plants resolved into one well-defined clade of helper NLRs (NRC-H) and two sister clades comprising the NRC-dependent sensor NLRs (NRC-S) (Fig. 1C).

**Figure 1.**
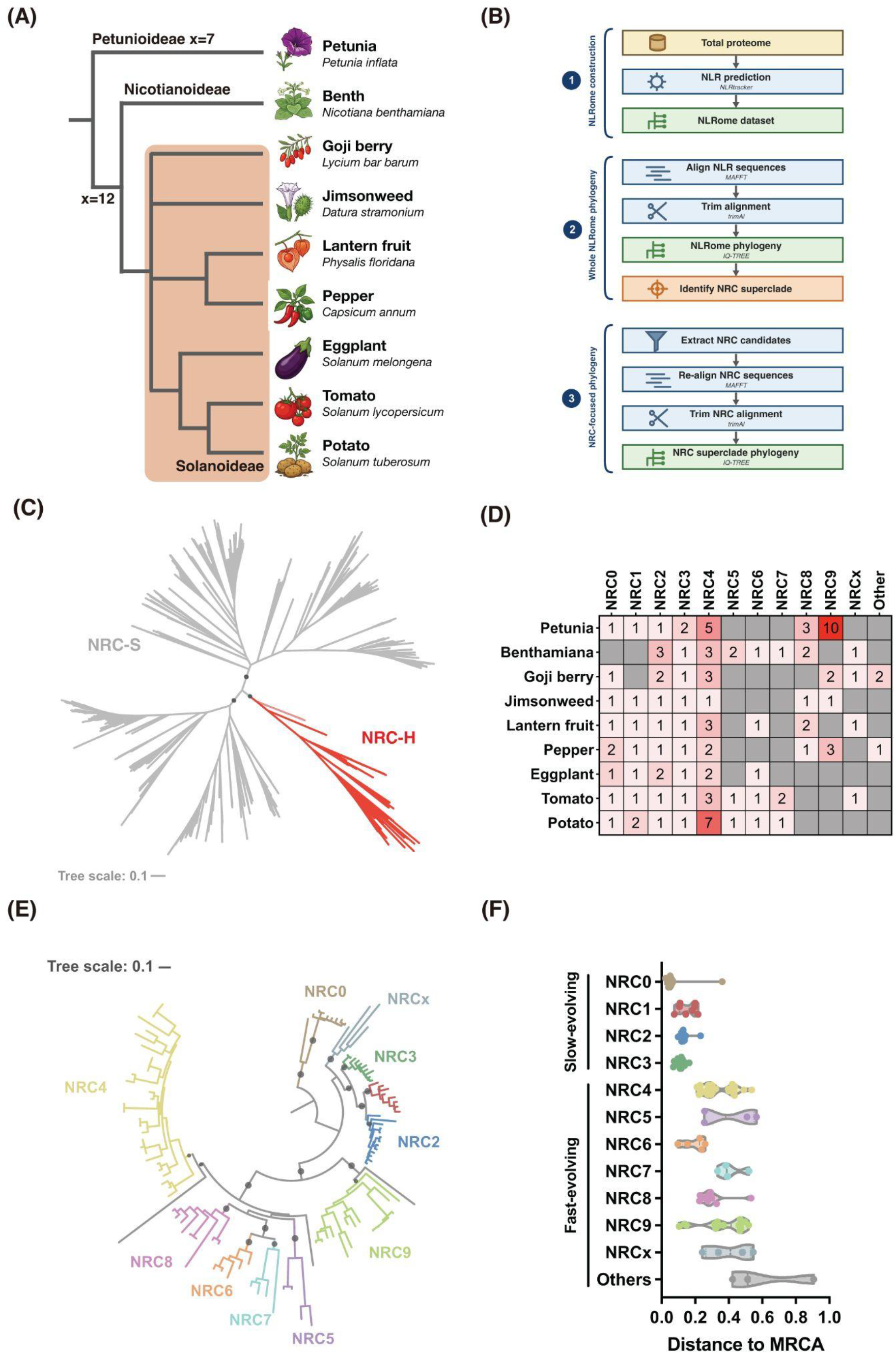
Phylogenetic distribution and diversification of the NRC immune receptor network across solanaceous plants. (A) Simplified phylogenetic relationships among representative solanaceous species included in this study. (B) Workflow for identification and phylogenetic classification of NRC helper NLRs. (C) Phylogenetic reconstruction of the NRC superclade resolved one well-defined clade of helper NLRs (NRC-H; red) and a larger clade of NRC-dependent sensor NLRs (NRC-S; gray). Major branches with bootstrap values over 70 are marked with grey dots (D) Distribution and copy number variation of NRC helper subfamilies across different solanaceous species. Numbers indicate the number of genes identified in each NRC subgroup, highlighting lineage-specific expansion and diversification within the NRC network. (E) Phylogenetic relationships among NRC helper NLRs across solanaceous species. NRC subgroups are highlighted with different colors, and branches with bootstrap values over 70 are marked with grey dots. (F) The relative evolutionary divergence from the most recent common ancestor (MRCA), illustrating both conserved and rapidly evolving NRC lineages.

We next examined the detail of the NRC-H subclade and identified 11 distinct NRC helper subfamilies (Fig. 1D). Among these, NRC2, NRC3, and NRC4 were the only three subfamilies present across all nine Solanaceae species, consistent with their central roles in the canonical NRC network, as most known NRC-dependent sensor NLRs signal through these three NRC subfamilies (Wu et al. 2017; Witek et al. 2021; Lin et al. 2022). In contrast, the remaining NRC subfamilies displayed clear presence/absence polymorphisms across the nine species (Fig. 1D). NRC5 and NRC7 were identified only in *N. benthamiana* and the *Solanum* species (eggplant, tomato and potato), whereas NRC6 was present only in *N. benthamiana*, lantern fruit, and the *Solanum* spp. NRC8 and NRC9 were found only in species that fall outside the *Solanum* lineage (Fig. 1D), suggesting these variants may have been lost in ancestral *Solanum*. In each species, most NRC subfamilies contained only one to two homologs; however, the NRC4 subfamily exhibited duplications in nearly all species examined (Fig. 1D). Notably, petunia possessed an exceptional multiplication of 10 NRC9 homologs, representing a striking lineage-specific expansion event (Fig. 1D).

We then reconstructed the phylogeny of all solanaceous NRC variants and quantified branch lengths relative to their most recent common ancestor (MRCA). This analysis revealed striking heterogeneity in the extent of sequence divergence among NRC subfamilies. NRC0, NRC1, NRC2, and NRC3 exhibited short branch lengths (<0.2), indicating relatively slow rates of sequence divergence and likely strong evolutionary constraint (Figs. 1E and 1F, Datasets S2-S4). In contrast, NRC4, NRC5, NRC6, NRC7, NRC8, NRC9, and NRCx displayed substantially longer branch distances (>0.2), reflecting rapid diversification (Figs. 1E and 1F). These patterns suggest that while a core set of NRC helpers remained highly conserved across Solanaceae, other NRC subfamilies diversified more rapidly and may have contributed to lineage- or species-specific immune functions.

### The NRC-dependent sensor NLR repertoire exhibits extensive lineage-specific expansion across Solanaceae

Next, we examined the diversification of NRC-dependent sensor NLRs across nine representative solanaceous species. Maximum-likelihood phylogenetic analysis resolved these receptors into 14 major subfamilies, including previously defined CNL-1, CNL-2, CNL-3, CNL-9 to CNL-13, NRC0-S and newly defined CNL-17 to CNL-21 (Fig. 2A, Datasets S5-S7). Consistent with previous studies, functionally characterized resistance genes mapped to their expected subfamilies, including Rpi-blb2 and Mi-1.2 (CNL-1), Rx/Rx2 (CNL-2), Rpi-amr1 (CNL-3), Hero (CNL-9), Sw5b (CNL-10), R1 and Prf (CNL-11), Rpi-amr3 (CNL-13), and Bs2 (CNL-17), providing further support for the inferred phylogeny (Andolfo et al. 2014; Seo et al. 2016; Witek et al. 2021; Sakai et al. 2024).

**Figure 2.**
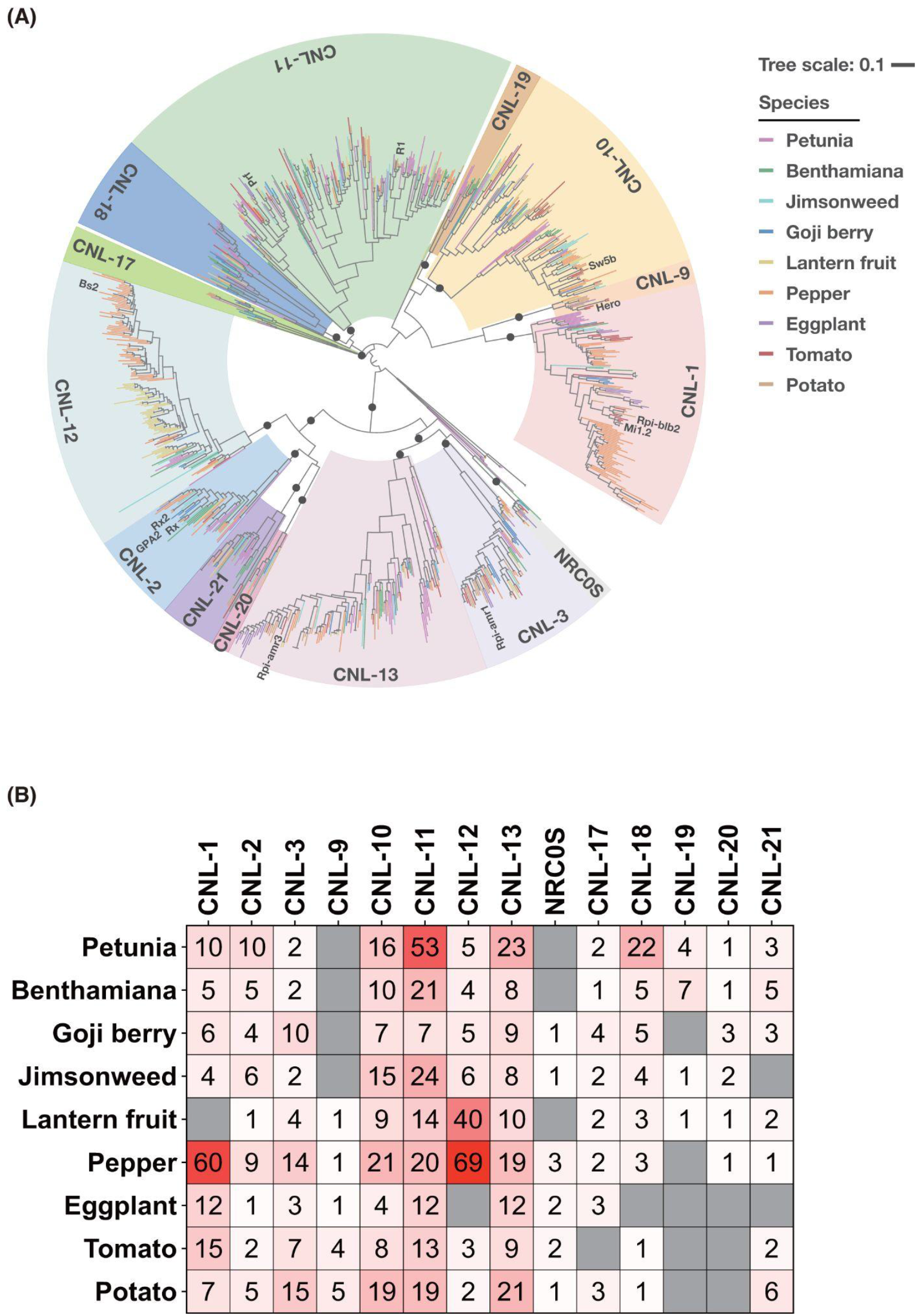
Phylogenetic classification of NRC-dependent sensor NLRs across nine representative solanaceous species. (A) Maximum-likelihood phylogenetic tree of NRC-dependent sensor NLR proteins identified from nine representative solanaceous species. Branches are color-coded according to species. CNL subfamilies are indicated by the shaded sectors surrounding the tree. Representative functionally characterized NLRs are labeled to illustrate the phylogenetic placement of known immune receptors. Black circles indicate nodes with strong bootstrap support (≥70%). Scale bar represents 0.1 amino acid substitutions per site. (B) Size of NRC-dependent sensor NLR subfamilies across the nine solanaceous species. Numbers indicate the number of proteins assigned to each subfamily in each species. Color intensity reflects gene copy number, with darker red indicating larger family sizes. Gray boxes indicate the absence of members in the corresponding subfamily.

Although all species retained representatives of most CNL subfamilies, their sizes differed markedly (Fig. 2B). Several subfamilies, including CNL-2, CNL-3, CNL-9, NRC0s, CNL-17, CNL-19, CNL-20, and CNL-21, remained relatively small across species. In contrast, others underwent substantial lineage-specific expansion. Pepper showed dramatic expansion of CNL-1 (60 genes) and CNL-12 (69 genes), while lantern fruit also possessed an expanded CNL-12 repertoire (40 genes). Petunia exhibited the largest expansion of CNL-11 (53 genes), together with relatively large CNL-13 and CNL-18 subfamilies. These results demonstrate that, despite a conserved overall phylogenetic framework, the non-helper CNL repertoire has diversified extensively through lineage-specific expansion and contraction during Solanaceae evolution.

### Dynamic evolution of NRC helper–sensor genomic clusters across Solanaceae

Our previous analyses suggest that the NRC network originated from an ancestral helper– sensor NLR gene cluster resembling NRC0 and its linked sensor NLRs (Goh et al. 2024; Sakai et al. 2024). To assess the extent to which solanaceous NRC helpers remain organized in genomic clusters with putative sensor NLRs, we quantified their chromosomal proximity based on previously established criteria, defining genes located within 50 kb of one another as belonging to the same cluster (Sakai et al. 2024). Using these criteria, helper– sensor NLR clusters (H-S) and helper–helper–sensor NLR clusters (H-H-S) were identified in most species examined. A notable exception was *N. benthamiana*, in which none of the NRC genes were physically linked to sensor NLRs at their genomic loci. In tomato and petunia, many NRCs were found within either H-S or H-H-S clusters, whereas relatively few helper–sensor associations were detected in pepper and goji berry (Fig. S1A). Helper– helper NLR clusters lacking an associated sensor NLR (H-H) were also observed in most species, with tomato representing the only exception in which such arrangements were not detected (Fig. S1A).

Analysis of sensor NLR genomic organization revealed that sensor–sensor NLR clusters (S-S) are particularly abundant in tomato, potato, lantern fruit, and petunia, whereas eggplant, goji berry, jimsonweed, and *N. benthamiana* harbor substantially fewer such clusters (Fig. S1B). In contrast to helper NLRs, only a small proportion of sensor NLRs were associated with helper NLRs, either in sensor–helper clusters (S-H) or in sensor–sensor–helper clusters (S-S-H). This pattern is consistent with the hypothesis that, following the ancestral loss of physical linkage between helper and sensor NLRs, the two classes largely evolved independently and subsequently followed distinct evolutionary trajectories (Goh et al. 2024; Sakai et al. 2024).

Several helper–sensor genomic clusters were shared across solanaceous plants of different genus, whereas corresponding linkages appear to have been lost in others. As the most ancient NRC variant (Sakai et al. 2024), NRC0 in goji berry, jimsonweed, and the *Solanum* species cluster with their putative matching sensor NLRs, which also grouped together phylogenetically (Fig. 3A). However, this NRC0–sensor linkage was absent in other examined species, suggesting that the ancestral genomic association was lost independently during Solanaceae evolution. Furthermore, while NRC6 orthologs were identified in *N. benthamiana,* lantern fruit, and the *Solanum* species, the reported linkage to Hero homologs within the CNL-9 clade was detected only in lantern fruit and the *Solanum* species, but not in *N. benthamiana* (Fig. 3B) (Lüdke et al. 2025). Similarly, NRC8 variants in lantern fruit/pepper and jimsonweed also formed genomic clusters with sensor NLRs belonging to the CNL-10 and CNL-12, respectively (Fig. 3B); however, whether these sensors functionally depend on NRC8 remains to be determined. Besides the conserved helper–sensor NLR clusters, helper–helper associations between NRC3 and NRC9 were also observed, forming clustered loci in petunia, goji berry, jimsonweed and pepper species (Fig. 3C). Because these genomic associations were conserved across species from different genera, we propose that these clusters originated early during Solanaceae evolution and were subsequently lost in multiple lineages.

**Figure 3.**
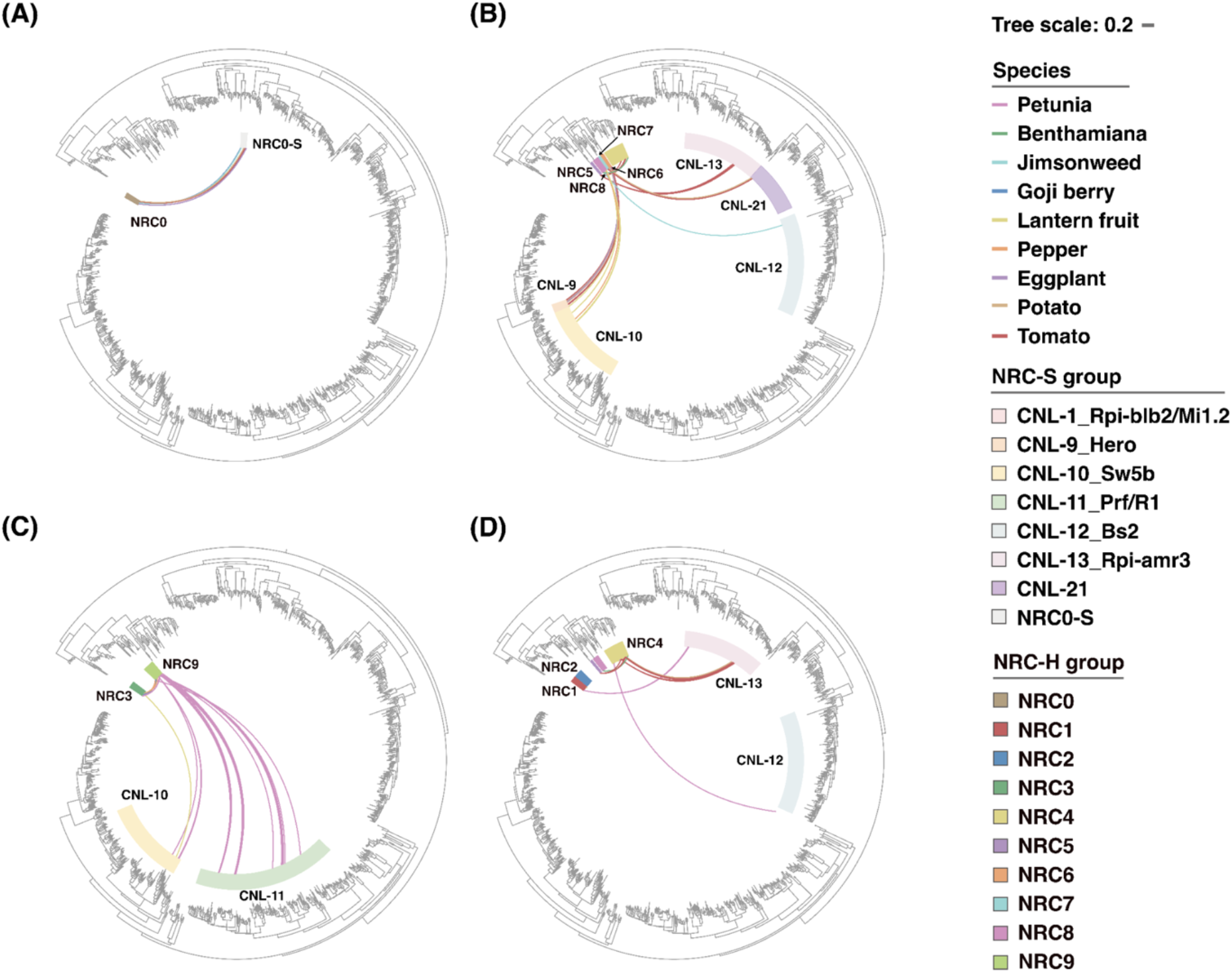
Genomic architecture of helper–sensor NLR clusters across Solanaceae. Genomic clusters containing NRC helper and sensor NLRs were identified from gene model annotations of nine solanaceous species. Genes located within 50 kb of each other were considered part of the same genomic cluster. Colored links connect helper and sensor NLR subfamilies found within the same genomic clusters, and branch colors indicate species. (A) NRC0–sensor clusters identified in jimsonweed, goji berry, and *Solanum* species, with associated sensor NLRs belonging to the NRC0S CNL group. (B) NRC5-, NRC6-, NRC7-, and NRC8–sensor clusters identified in *Solanum* species and lantern fruit, with associated sensor NLRs predominantly belonging to the CNL-9 (Hero) and CNL-13 (Rpi-amr3) groups. (C) Extensive NRC9–sensor clustering in petunia, with associated sensor NLRs mainly belonging to the CNL-11 (Prf/R1/9S) group. (D) Additional helper–sensor clusters involving NRC1 and NRC4 across different solanaceous species, with associated sensor NLRs belonging primarily to the CNL-13 (Rpi-amr3) group.

In addition to the intergeneric conserved helper–sensor clusters described above, several helper–sensor associations exhibited clear lineage- or species-specific patterns, indicating a more dynamic evolutionary history. Among these, the most striking example was observed for NRC9. In petunia, NRC9 formed extensive genomic clusters with multiple sensor NLRs, including members of the CNL-10 and CNL-11 clades (Fig. 3C). In contrast, no comparable NRC9–sensor clusters were detected in goji berry, jimsonweed, or pepper, despite the presence of NRC9 homologs in these species. A similar pattern was observed for NRC1. Although NRC1 homologs were retained across most examined species, only the petunia NRC1 locus was associated with a nearby sensor NLR belonging to the CNL-13 clade, for which the functional relationship remains unknown (Fig. 3D). Likewise, NRC3 was conserved across all species analyzed, yet helper–sensor linkage was detected exclusively in lantern fruit (Fig. 3C).

NRC4-associated clusters also showed substantial variation among species. NRC4 variants in petunia, potato, and tomato were found within helper–sensor clusters. In potato and tomato, the associated sensors belonged to the CNL-13 group, whereas the linked sensor in petunia does not belong to any defined NLR clade (Fig. 3D). The potato and tomato NRC4 clusters also contained NRC5; however, NRC5 in *N. benthamiana* was not associated with this cluster. While NRC7 in *Solanum* species formed clusters with putative sensor NLRs belonging to the CNL-13 group, no corresponding NRC7-associated cluster was detected in *N. benthamiana* (Fig. 3B). Nevertheless, the *N. benthamiana* NRC7 locus was located in proximity to two sensor NLRs belonging to the CNL-18 and CNL-21 groups, although both associations exceeded the 50-kb cutoff used to define NLR clusters (Fig. S2). However, their functional relationship has not been tested. Together, the patchy distribution of these clusters across closely related species suggests that many helper–sensor associations arose relatively recently or were independently lost during evolution. In contrast, NRC2 stood out among all NRC helper subfamilies in that no genomic association with sensor NLRs was detected in any examined species, suggesting that the loss of helper–sensor linkage occurred early during NRC2 evolution.

### A cross-Solanaceae functional atlas of NRC helper activity reveals conserved cores and lineage-specific diversification

Our previous work on NRC3 revealed that diversification among NRC orthologs drives subfunctionalization, thereby reshaping the genetic architecture of NRC immune networks (Huang et al. 2024). To more comprehensively test how evolutionary diversification has influenced helper–sensor compatibility across Solanaceae, we cloned NRC variants from representative species and systematically assessed their ability to complement sensor NLR– mediated cell death in *NRC2/3/4* triple-knockout *N. benthamiana* (*nrc2/3/4*) that is also missing NRC0, NRC1 and NRC9 (Fig. 1A) (Wu et al. 2020). For clarity, we use the prefixes Pe, Nb, Lb, Ds, Pf, Ca, and Sl to denote NRC variants from petunia, *N. benthamiana*, goji berry, jimsonweed, lantern fruit, pepper, and tomato, respectively. For sensor NLRs, we included previously defined *NRC2/3/4*-dependent (Rx, Bs2, Rpi-amr3, Sw5), *NRC2/3*-dependent (Rpi-amr1 and Prf (Pto/AvrPto)), *NRC4*-dependent (Rpi-blb2 and R1), and *NRC6*-dependent (HeroF) sensors, as well as a newly cloned petunia sensor genetically linked to PeNRC9, named PeNRC9-S.

Among the four NRC1 variants tested, we did not detect activity for DsNRC1, whereas PeNRC1, CaNRC1, and SlNRC1 were able to mediate cell death induced by Rx, Rpi-amr3, and Bs2 (Fig. 4A; Figs. S3-S5). Interestingly, only PeNRC1 also functioned with Prf and Rpi-amr1, whereas the other NRC1 variants did not (Fig. 4A; Figs. S6 and S7). This pattern suggests either a gain of compatibility in PeNRC1 with Rpi-amr1/Prf or a loss of functional interaction between these sensors and the Ca/SlNRC1 variants.

**Figure 4.**
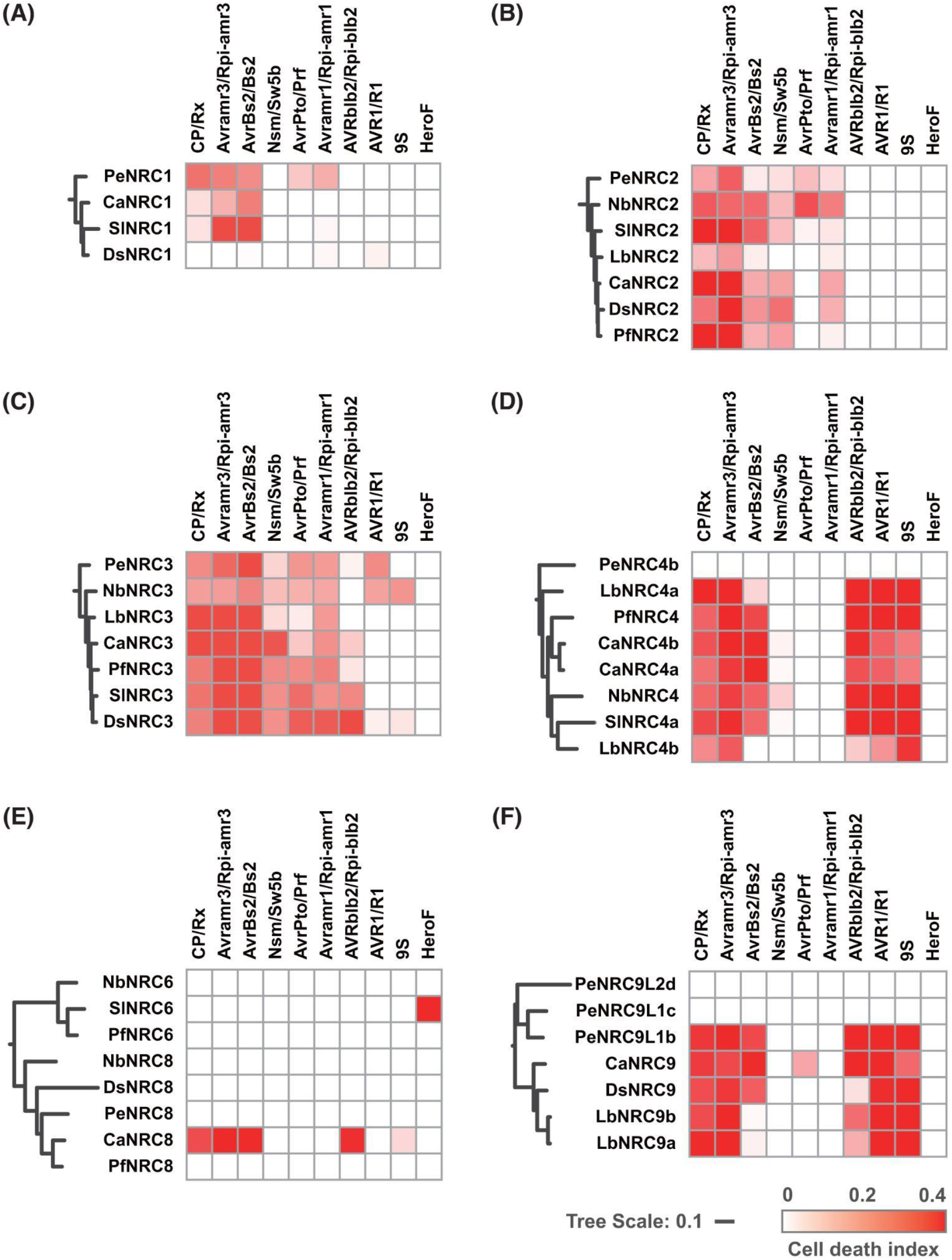
Functional compatibility matrix of solanaceous NRC helpers with diverse effector–sensor NLR pairs. Cell death assays were conducted in *NRC2/3/4* triple-knockout *N. benthamiana* triggered by pairing each NRC helper variant from seven Solanaceae species (petunia, *N. benthamiana*, goji berry, jimsonweed, lantern fruit, pepper, and tomato) with a panel of canonical effector–sensor NLR combinations. Tested pairs include CP/Rx, Avramr3/Rpi-amr3, AvrBs2/Bs2, Nsm/Sw5b, AvrPto/Prf, Avramr1/Rpi-amr1, AVRblb2/Rpi-blb2, AVR1/R1, and two autoactive sensors (petunia 9S and tomato HeroF). Heatmaps summarize the degree of cell death responses across helper–sensor combinations involving (A) NRC1, (B) NRC2, (C) NRC3, (D) NRC4, (E) NRC6/8, and (F) NRC9. Robust cell death, weak cell death, and no response are indicated by graded color intensities. The phylogenetic trees of NRC helpers were extracted from the master NRC helper tree shown in Fig. 1E.

Across the seven NRC2 variants tested, all retained the ability to function with Rx and Rpi-amr3. However, several loss-of-compatibility events were observed: PeNRC2 did not function effectively with Bs2 (Fig. 4B; Fig. S5); SlNRC2 failed to mediate Prf cell death and had weak activity with Rpi-amr1 (Fig. 4B; Figs. S6 and S7); LbNRC2 functioned only with Rx and Rpi-amr3 but not efficiently with other sensor NLRs tested (Fig. 4B; Figs. S3 and S4); Many NRC2 homologs, including LbNRC2, CaNRC2, DsNRC2, and PfNRC2, failed to effectively function with Prf (Fig. 4B; Fig. S6), suggesting that Prf compatibility may have been lost in the common ancestor of this NRC2 clade, although the observed pattern could also reflect incompatibility with the endogenous Prf signaling context in *N. benthamiana*.

In contrast to NRC2, the seven NRC3 orthologs tested appear to be more functionally conserved, with all variants supporting cell death mediated by Rx, Rpi-amr3, Bs2, Prf, Rpi-amr1 and Sw5b (Fig. 4C; Figs. S3-S8). However, PeNRC3, NbNRC3 and LbNRC3 did not function with Rpi-blb2, (Fig. 4C; Fig. S9) consistent with the previously reported loss-of-function mutation in these variants (Huang et al. 2024). Interestingly, PeNRC3 and NbNRC3 are the only two NRC variants able to function with R1 in complementation assays (Fig. 4C; Fig. S10), despite R1 being characterized as NRC4-dependent, suggesting an ancestral loss of compatibility with R1 in other NRC3 lineages. Furthermore, two distantly related NRC3 variants, NbNRC3 and DsNRC3, were able to function, albeit weakly, with R1 and the petunia sensor PeNRC9-S (Fig. 4C; Fig. S11), suggesting independent gains of compatibility in these orthologs.

Although NRC4 is highly divergent and rapidly evolving (Fig. 1), its functional activity is surprisingly well conserved. With the exception of PeNRC4b, which did not exhibit detectable cell death with any tested sensor NLR, all other NRC4 variants tested were able to induce cell death in response to Rx, Rpi-amr3, Bs2, Rpi-blb2, R1, and PeNRC9-S (Fig. 4D; Fig. S3-5 and S9-S11). Interestingly, while LbNRC4a retained compatibility with Bs2, LbNRC4b failed to do so (Fig. 4D), indicating that paralogs within the same species can evolve distinct sensor compatibility. Notably, NbNRC4 was the only NRC4 variant capable of rescuing Sw5b-mediated cell death (Fig. 4D; Fig. S8), raising the possibility that Sw5b may be functionally NRC2/3-dependent, rather than strictly NRC2/3/4-dependent, when operating outside of *Nicotiana*.

The cell death responses mediated by NRC6 and NRC8 across different sensor NLR combinations displayed a highly restricted pattern, with most pairings showing little to no activity. Among NRC6 variants, only SlNRC6 triggered a robust cell death response when paired with HeroF (Fig. 4E; Fig. S12), indicating a narrow and sensor-specific compatibility. Interestingly, within the NRC8 group, CaNRC8 was the only tested variant capable of inducing cell death, responding to a subset of sensors including Rx, Rpi-amr3, Bs2, Rpi-blb2, and PeNRC9-S (Fig. 4E; Figs. S3-5, S9 and S11). In contrast, other NRC8 variants tested failed to elicit detectable responses across all tested combinations.

The cell death responses mediated by NRC9 were strikingly similar to those observed for NRC4. With the exception of PeNRC9L2d and PeNRC9L1c, which failed to induce detectable cell death with any tested sensor NLR, all NRC9 variants effectively mediated cell death triggered by Rx, Rpi-amr3, Rpi-blb2, R1, and PeNRC9-S (Fig. 4F; Figs. S3, S4, and S9-S11). In addition, most NRC9 variants—except for LbNRC9a and LbNRC9b—functioned with Bs2 to induce cell death (Fig. 4F; Fig. S5). Only CaNRC9 supported Prf-mediated cell death, highlighting a unique compatibility within this lineage (Fig. 4F; Fig. S6). These results indicate that NRC9 broadly retains robust helper activity across multiple sensor NLRs, closely resembling the functional profile of NRC4 while the observed exceptions suggest fine-scale lineage-specific divergence in compatibility.

To evaluate whether the lack of activity reflected signaling incompetence, we generated autoactive variants of nonresponsive helpers (PeNRC4b, PfNRC6, PeNRC8, NbNRC8, DsNRC8, and PfNRC8). Among these, only PfNRC6DV induced a clear cell death phenotype (Fig. S13C). The remaining “autoactive” helper variants did not produce cell death phenotypes (Fig. S13), suggesting that additional mutations within these NRC variants may have resulted in loss of function.

### A divergent petunia NRC0 exhibits different compatibility with NRC0-dependent sensors compared to other solanaceous NRC0

We previously reported that NRC0 is conserved across asterid lineages and functions with a unique group of sensor NLRs that are often tightly linked on the chromosome. Consistent with this, NRC0 variants did not trigger detectable cell death with any of the tested effector– sensor combinations (Fig. S14) (Goh et al. 2024; Sakai et al. 2024). Interestingly, we noticed that the petunia NRC0 variant (PeNRC0) did not cluster with other solanaceous NRC0 proteins and instead displayed a long branch in the phylogenetic tree, suggesting that it may represent a distinct early evolved NRC0 lineage (Fig. 1). To further investigate this, we performed a phylogenetic analysis including NRC0 homologs from additional asterid species. This analysis revealed that PeNRC0 forms a separate clade and does not group with either the previously described NRC0 lineage or the Asterales- and Ericales-specific NRC0 subclades (Fig. 5A, Datasets S8 and S9). This pattern likely reflects an early divergence of PeNRC0 in the ancestral petunia lineage, while the ancestral NRC0 has been lost.

**Figure 5.**
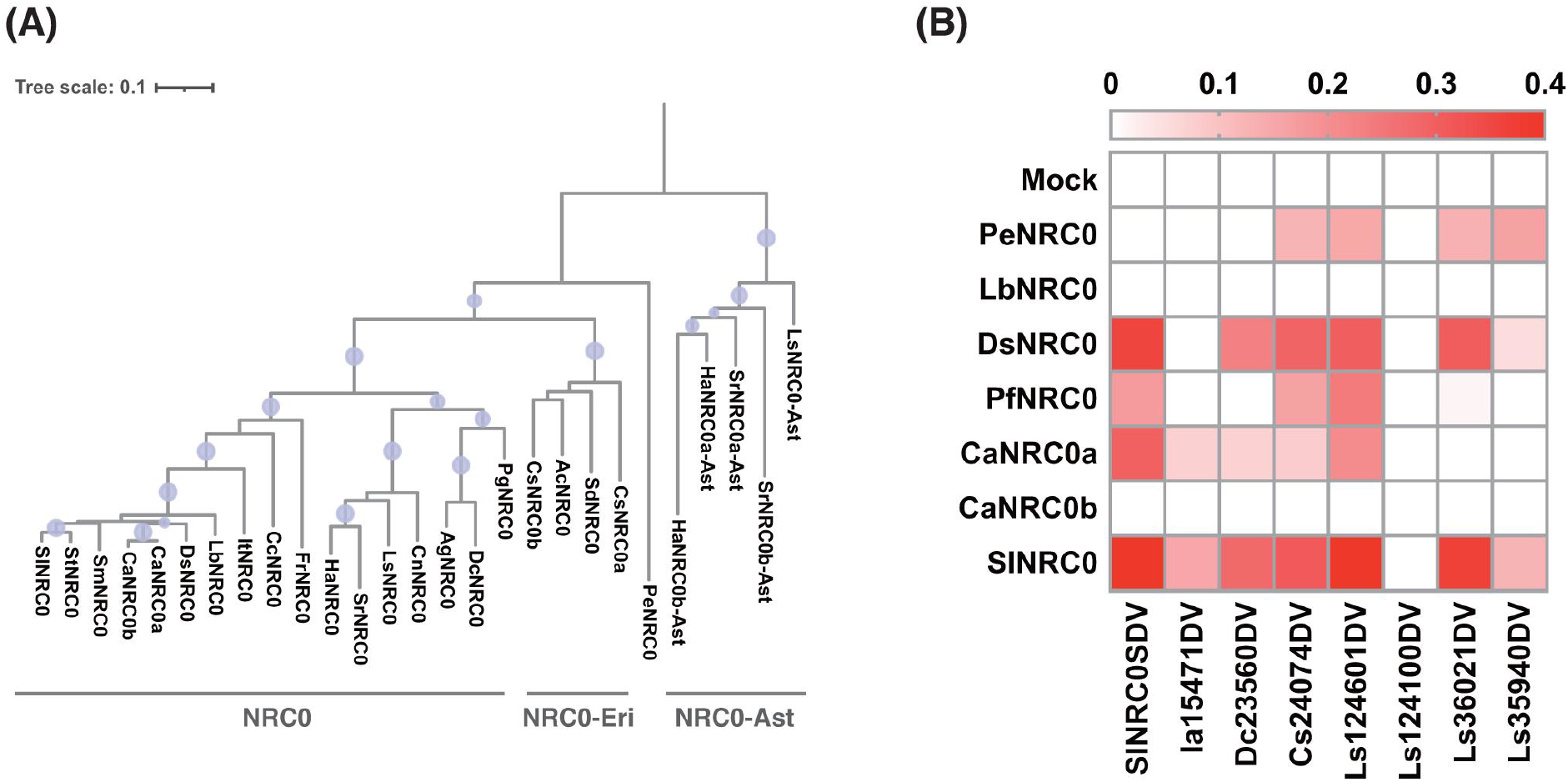
Phylogenetic and functional divergence of NRC0 variants across asterid species. (A) Maximum-likelihood phylogeny of NRC0 helper NLRs from Solanaceae and representative asterid species. Branches with bootstrap values over 70 are marked with purple circles. (B) Functional profiling of NRC0 variants using autoactive NRC0-dependent sensor NLRs from *Solanum lycopersicum*, *Ipomoea aquatica*, *Daucus carota*, *Camellia sinensis*, and *Lactuca sativa*. Robust cell death, weak cell death, and no response are indicated by graded color intensities.

To determine whether this phylogenetic divergence is consistent with functional diversification, we tested a panel of autoactive NRC0-dependent sensor NLRs with PeNRC0 and other NRC0 variants from solanaceous plants. Consistent with previous reports, SlNRC0 supported robust cell death responses with most sensor NLRs, with the exception of Ls124100DV (Fig. 5B; Figs. S15A–S15H) (Goh et al. 2024). Similar overall compatibility patterns were observed for DsNRC0, PfNRC0, and CaNRC0a, although each displayed distinct incompatibilities, failing to function with some of the tested NRC0-dependent sensor NLRs (Fig. 5B; Figs. S15A–S15H). In contrast, LbNRC0 and CaNRC0b were completely non-responsive in all tested combinations (Fig. 5B; Figs. S15A–S15H).

Interestingly, PeNRC0 displayed a distinct compatibility profile, supporting cell death only with several sensor NLRs from *C. sinensis* and *L. sativa*, while failing to function with SlNRC0-SDV, a solanaceous sensor NLR that was compatible with most other solanaceous NRC0 variants tested (Fig. 5B; Figs. S15D, S15E, S15G, S15H). This compatibility profile differed markedly from those of other solanaceous NRC0 proteins tested in this study, but more similar to the CsNRC0 belonging to NRC0-Eri reported previously (Goh et al. 2024). Together, these findings support the hypothesis that PeNRC0 represents an early-diverging NRC0 lineage that is functionally distinct from previously characterized NRC0 clades.

## Discussion

The NRC immune receptor network represents one of the best-characterized examples of a complex helper–sensor NLR system in plants. Previous studies established the existence of NRC-dependent signaling pathways in diverse lineages of asterids including Solanaceae (Goh et al. 2024; Sakai et al. 2024). However, in Solanaceae, the analysis was limited to species in the genera *Nicotiana* and *Solanum* (Wu et al. 2017; Witek et al. 2021; Lin et al. 2022). Here, by integrating comparative phylogenomics and systematic cross-species functional assays across nine solanaceous species, we reveal that the NRC network exhibits a striking combination of deep conservation and extensive lineage-specific diversification. Our analyses identify both conserved core helper lineages that remain broadly interchangeable across Solanaceae and rapidly evolving peripheral NRC variants associated with more restricted or lineage-specific compatibility profiles (Fig. 6). Together, these findings provide a broader evolutionary framework for understanding how helper–sensor immune networks diversify while maintaining functional robustness.

**Figure 6.**
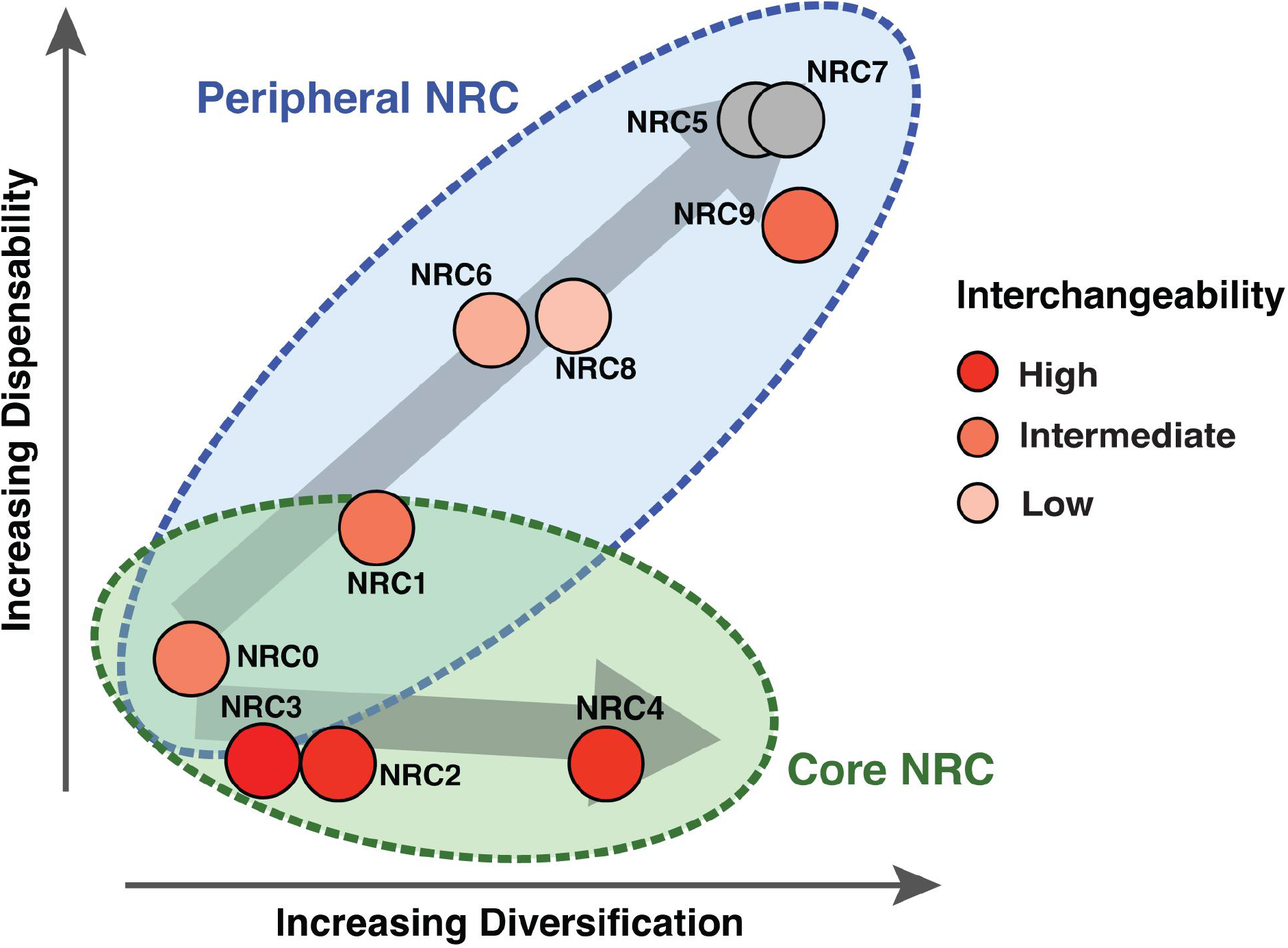
Evolutionary diversification and functional interchangeability of NRC helper NLRs across Solanaceae. Conceptual model summarizing the evolutionary and functional characteristics of NRC helper subfamilies. NRC helpers are positioned according to their relative degree of sequence diversification (x-axis) and dispensability across solanaceous species (y-axis). NRC2, NRC3, NRC4 constitute the classical core NRC group, which is broadly retained across Solanaceae, whereas NRC5, NRC6, NRC7, NRC8, and NRC9 represent peripheral NRCs that show greater lineage-specific variation and dispensability. NRC0 and NRC1 display intermediate characteristics and overlap with both the core and peripheral NRC groups. Arrows indicate the distinct evolutionary trajectories of the core and peripheral NRC groups. Circle colors indicate the degree of functional interchangeability among orthologs from different solanaceous species, ranging from high (red) to intermediate (orange) and low (light orange); gray indicates subfamilies for which interchangeability was not determined.

Our phylogenomic analyses resolved 11 NRC helper subfamilies with markedly different evolutionary patterns. NRC2, NRC3, and NRC4 were conserved across all examined solanaceous species, consistent with their central role in supporting a broad range of sensor NLRs (Wu et al. 2017). In contrast, other NRC lineages displayed different degrees of presence/absence polymorphisms and lineage-specific expansions, including the remarkable amplification of NRC9 in petunia (Fig. 1D). These observations suggest that the NRC network consists of a conserved signaling core and a dynamic periphery shaped by repeated duplication, diversification, and lineage-specific gene loss during Solanaceae evolution (Fig. 6).

Despite extensive diversification in sequences and genomic organization, many homologous NRC variants retained surprisingly broad functional interchangeability. In particular, NRC2, NRC3, NRC4, and NRC9 variants from distantly related species frequently showed functional interchangeability, working with the same set of sensor NLRs in cell death assays in *N. benthamiana* (Fig. 4). This was especially striking for NRC4 and NRC9, which exhibited relatively long phylogenetic branch lengths yet maintained highly conserved helper activity across multiple sensor combinations (Fig. 1E). In contrast, closely related orthologs with short phylogenetic distances occasionally displayed distinct compatibility profiles (Fig. 4). For example, our previous study on NRC3 identified two amino acid substitutions that altered compatibility with specific sensor NLRs despite the overall high sequence similarity among NRC3 orthologs (Huang et al. 2024). Similar patterns were also observed among NRC0, NRC1, and NRC2 orthologs, in which closely related variants differed in their ability to function with particular sensor NLRs (Fig. 4). Together, these findings indicate that phylogenetic distance alone is not a strong predictor of sensor–helper compatibility. Instead, helper–sensor signaling competence appears robust to substantial evolutionary divergence, whereas relatively small sequence changes can be sufficient to alter compatibility with individual sensor NLRs. Interestingly, recent AlphaFold3 predictions of the Rx–NRC2 complex have begun to reveal the molecular interface between sensor and helper NLRs, providing new insights into the activation-and-release mechanism that underpins NRC sensor–helper signaling (Contreras et al. 2023b, 2024; Toghani et al. 2026a). These advances offer an attractive framework for interpreting the compatibility shifts observed among NRC variants and raise the possibility of engineering sensor–helper specificity (Toghani et al. 2026a). Future studies integrating structural predictions with experimental validation will be essential to determine whether the molecular interfaces identified in silico can broadly explain the patterns of compatibility and incompatibility observed across the NRC network of various plant species.

Our analyses of helper–sensor genomic clustering further support a highly dynamic evolutionary history of the NRC network. Several conserved helper–sensor associations, including NRC0-linked NLR clusters, were maintained across distantly related solanaceous genera, suggesting that these associations originated early during Solanaceae evolution (Fig. 3). In contrast, many other helper–sensor linkages were lineage-specific or inconsistently retained even among closely related species, indicating repeated independent losses or more recent evolutionary assembly. Importantly, the widespread absence of physical linkage between helper and sensor NLRs in many species suggests that genomic decoupling did not disrupt network functionality. Instead, helper and sensor NLRs appear capable of evolving semi-independently after the loss of physical association, potentially enabling flexible rewiring of compatibility relationships during evolution (Wu et al. 2017).

The functional analyses further revealed substantial asymmetry between helper lineages. NRC2, NRC3, NRC4, and NRC9 generally retained broad compatibility profiles, whereas NRC6 and NRC8 displayed highly restricted activities (Fig. 4). NRC0 variants were completely non-active with canonical NRC2/3/4-dependent sensors, consistent with the hypothesis that NRC0 represents a more specialized signaling module functioning with distinct linked sensor lineages (Goh et al. 2024; Sakai et al. 2024). Particularly intriguing was the identification of a divergent petunia NRC0 lineage with compatibility profiles distinct from canonical solanaceous NRC0 proteins (Fig. 5). The phylogenetic position and functional properties of PeNRC0 suggest that multiple ancient NRC0 lineages may have coexisted during early asterids evolution, followed by differential lineage-specific retention and loss.

Several observations also suggest that helper–sensor compatibility can evolve through both gain- and loss-of-function processes. For example, PeNRC1 displayed compatibility with Prf and Rpi-amr1 that was absent from closely related NRC1 homologs, whereas other NRC variants lost compatibility with otherwise conserved sensors (Fig. 4). Similarly, only two NRC3 variants retained compatibility with R1, despite broad conservation of NRC3 helper activity overall (Fig. 4C). These patterns suggest that helper–sensor relationships are evolutionarily plastic and may be repeatedly rewired through relatively limited molecular changes. Future structural and biochemical analyses will be necessary to determine how evolutionary changes in helper–sensor recognition and signaling contribute to these compatibility shifts.

Overall, our study establishes a cross-Solanaceae evolutionary and functional framework for the NRC immune receptor network. The NRC network combines highly conserved core helper functions with extensive lineage-specific diversification in genomic organization, repertoire composition, and sensor compatibility. Importantly, our results demonstrate that large-scale genome restructuring and helper sequence diversification do not necessarily predict loss of functional interchangeability. Instead, NRC helpers appear to retain robust signaling competence over long evolutionary timescales while permitting flexible rewiring of specific helper–sensor interactions. These properties likely contributed to the remarkable evolutionary success and diversification of the NRC immune receptor network across Solanaceae.

## Materials and Methods

### Plant growth conditions

The *nrc2/3/4* knockout *Nicotiana benthamiana* plants were used for the cell death analyses. Plants were grown in a walk-in growth chamber under a 10-h-dark/14-h-light regime. The temperature was set as 28°C, and the humidity was maintained between 45% and 65%.

### Plasmid construction

Clones used in this study were generated using Golden Gate assembly following the Modular Cloning (MoClo) system. Level 0 plasmids encoding most NRC helper and sensor NLRs were synthesized as MoClo-compatible modules in kanamycin- or chloramphenicol-resistant pUC57 vectors by Twist Bioscience (South San Francisco, CA, USA) or Omics Bio (New Taipei City, Taiwan). For cloning NRC variants from petunia, full-length coding sequences were amplified from a cDNA library of *Petunia hybrida* V26 plants. Sequences containing internal BsaI or BpiI restriction sites were domesticated to generate MoClo-compatible versions. All amplified fragments were cloned into pAGM9121 to generate Level 0 modules and subsequently assembled into the binary vector pICSL86977, flanked by the CaMV 35S promoter and OCS terminator. To generate autoactive NRC variants, an Asp-to-Val (D→V) substitution was introduced by site-directed mutagenesis. Curated versions of pepper NRC2 and NRC3 were generated by amplifying the 5′ and 3′ regions from the corresponding Level 0 clones to remove inserted fragments. For the curated jimsonweed NRC2, the 5′ region of goji berry NRC2 and the 3′ region of jimsonweed NRC2 were amplified and assembled accordingly. All resulting amplicons were cloned into pICSL86977. Detailed sequence information and primer lists are provided in Table S2 and Table S3.

### Phylogenetic analysis

The NLRomes of nine solanaceous species were extracted from their respective proteome datasets using NLRtracker (Kourelis et al. 2021). The NB-ARC domain of each identified NLR protein was then isolated and subjected to multiple sequence alignment using MAFFT with the G-INS-i algorithm (Kuraku et al. 2013; Katoh et al. 2019). The unalign level was set to 0.8, with the option “Leave gappy regions” enabled. The resulting alignment was trimmed using trimAl with the “gappyout” method (Capella-Gutiérrez et al. 2009). Sequences shorter than 200 amino acids were omitted from the following analysis. The trimmed alignment was then used for phylogenetic reconstruction with IQ-TREE 2 (Trifinopoulos et al. 2016), applying the JTT+F+R4 substitution model and 1,000 ultrafast bootstrap replicates. The resulting phylogenetic tree was visualized and annotated using iTOL (Letunic and Bork 2007). The database information is provided in Table S1.

### Gene clustering analysis

Genomic coordinates of each NRC helper and sensor NLR were retrieved from their respective gene model annotation files. Genes located within 50 kb of each other were defined as belonging to the same genomic cluster. Clustering results were visualized and annotated using iTOL (Letunic and Bork 2007). The database information is provided in Table S1.

### Agrobacterium-mediated transient expression

*Agrobacterium tumefaciens* strain GV3101/pMP90 was used for transient gene expression assays. Bacterial strains were revived from glycerol stocks and cultured on 523 agar plates supplemented with appropriate antibiotics. After overnight incubation at 28 °C, bacterial cells were harvested by centrifugation at 5,000 × g for 5 min at room temperature. Pellets were resuspended in MMA buffer (10 mM MgCl₂, 10 mM MES-KOH, 200 μM acetosyringone, pH 5.6). Each construct was adjusted to an OD_600_ of 0.1 in the final infiltration mixture. Agrobacterial suspensions were infiltrated into the fourth and fifth fully expanded leaves of 27-day-old plants using 1 mL needleless syringes. At 3 days post-agroinfiltration, infiltrated leaves were harvested for photographic documentation and autofluorescence quantification.

### Cell death quantification

Autofluorescence emitted from dead cells was quantified using a UVP ChemStudio Imaging System (Analytik Jena). Raw autofluorescence images were acquired using blue LED illumination for excitation and a FITC emission filter (513–557 nm). Cell death levels were quantified based on autofluorescence intensity, with the exposure time fixed at 20 s to ensure optimal signal acquisition. Mean signal intensity was calculated using VisionWorks v11.2 software by manually selecting infiltrated regions and subtracting the corresponding background signal. The resulting values were normalized to the maximum possible intensity (65,535) to generate relative cell death intensities.

### Statistical analysis

Box plots and heatmaps were generated using GraphPad Prism version 10. Multiple comparisons were performed using the Kruskal–Wallis test followed by Dunn’s post hoc test, with the P-value threshold set at 0.05. Groups labeled with asterisks indicate statistically significant differences.

## Acknowledgments

We thank Mark Youles (SynBio, The Sainsbury Laboratory, UK) for sharing plasmids for molecular cloning, AmirAli Toghani and Sophien Kamoun (The Sainsbury Laboratory, UK) for helpful discussion and suggestion about the research, Doil Choi (Seoul National University, Korea) and Soohyun Oh (University of Seoul, Korea) for sharing plasmid materials.

## Author contributions

LYH and CHW designed the research. LYH, MHA, IBOO, and JK conducted the experiments. LYH and CHW analyzed the data. LYH and CHW wrote the manuscript.

## Data and materials availability

All data are available in the main text or the supplementary materials.

---

**Supplemental Figure S1.**
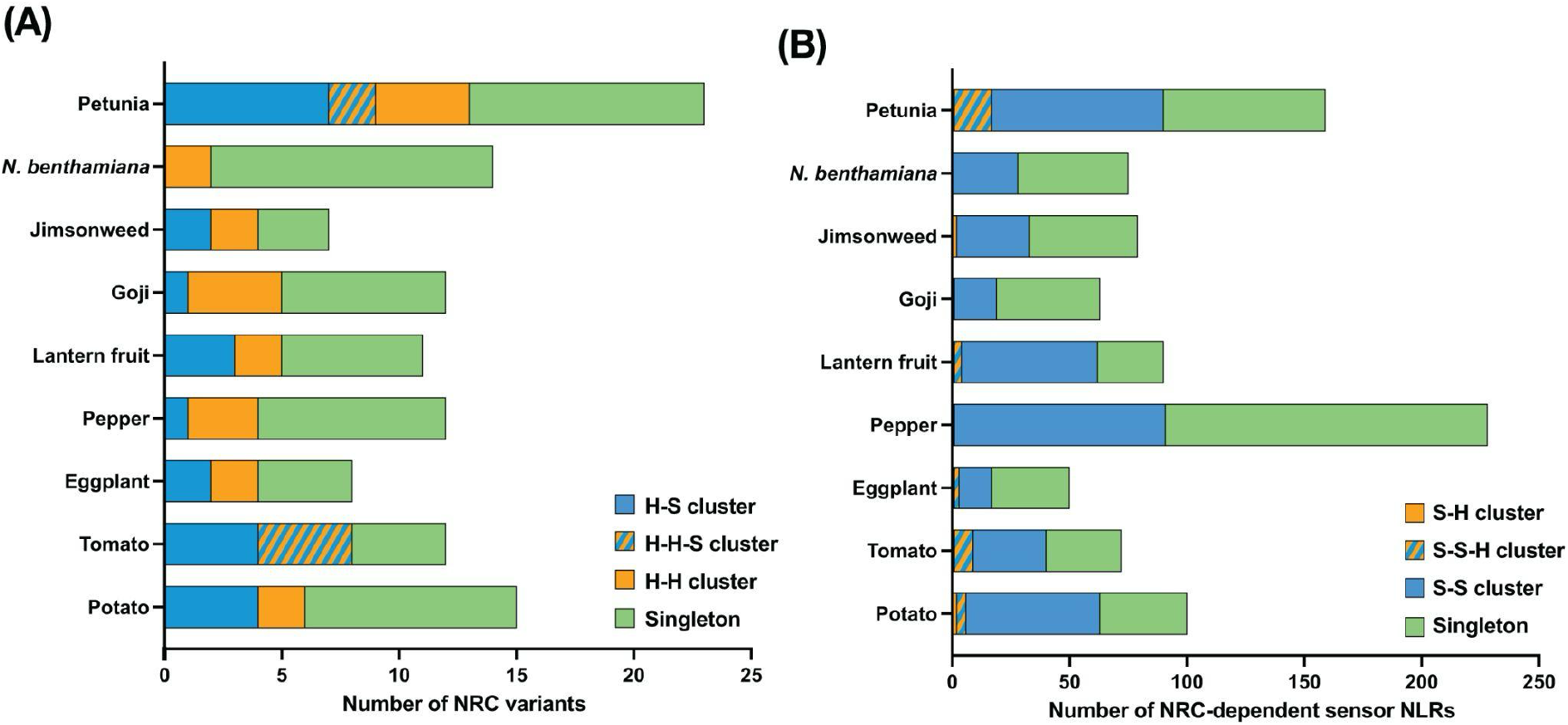
Distribution of helper–sensor and sensor–sensor NLR clusters across Solanaceae. (A) Number of NRC helper NLRs that are either clustered with other NRC helpers and/or NRC-dependent sensor NLRs or occur as isolated loci in each solanaceous species, based on a 50 kb genomic distance threshold for defining clusters. Genes clusters were classified as helper–sensor (H-S) clusters, helper–helper– sensor (H-H-S) clusters, helper–helper (H-H) clusters, or singletons. (B) Number of NRC-dependent sensor NLRs that are either clustered with other sensors or occur as isolated loci, using the same 50 kb clustering criterion, across the analyzed solanaceous genomes. Genes clusters were classified as sensor–helper (S-H) clusters, sensor–sensor–helper (S-S-H) clusters, sensor–sensor (S-S) clusters, or singletons.

**Supplemental Figure S2.**
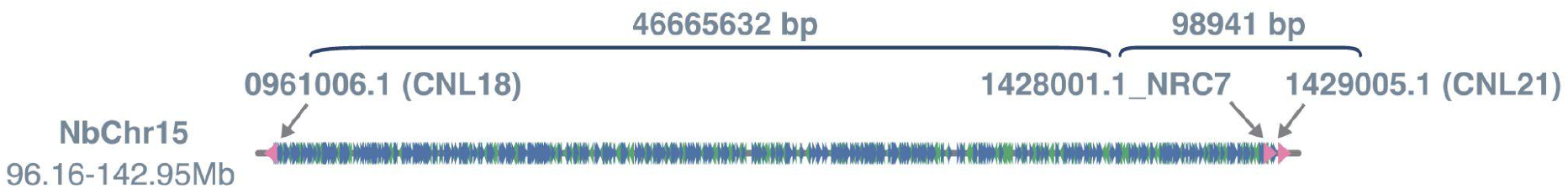
Genomic organization of the NRC7 locus in *Nicotiana benthamiana*. NbNRC7 (1428001.1_NRC7) is located on chromosome 15 (NbChr15) in proximity to two sensor NLRs belonging to the CNL-18 (0961006.1) and CNL-21 (1429005.1) subfamilies. The genomic distances between CNL-18 and NbNRC7 and between NbNRC7 and CNL-21 are approximately 46.7 Mb and 98.9 kb, respectively.

**Supplemental Figure S3.**
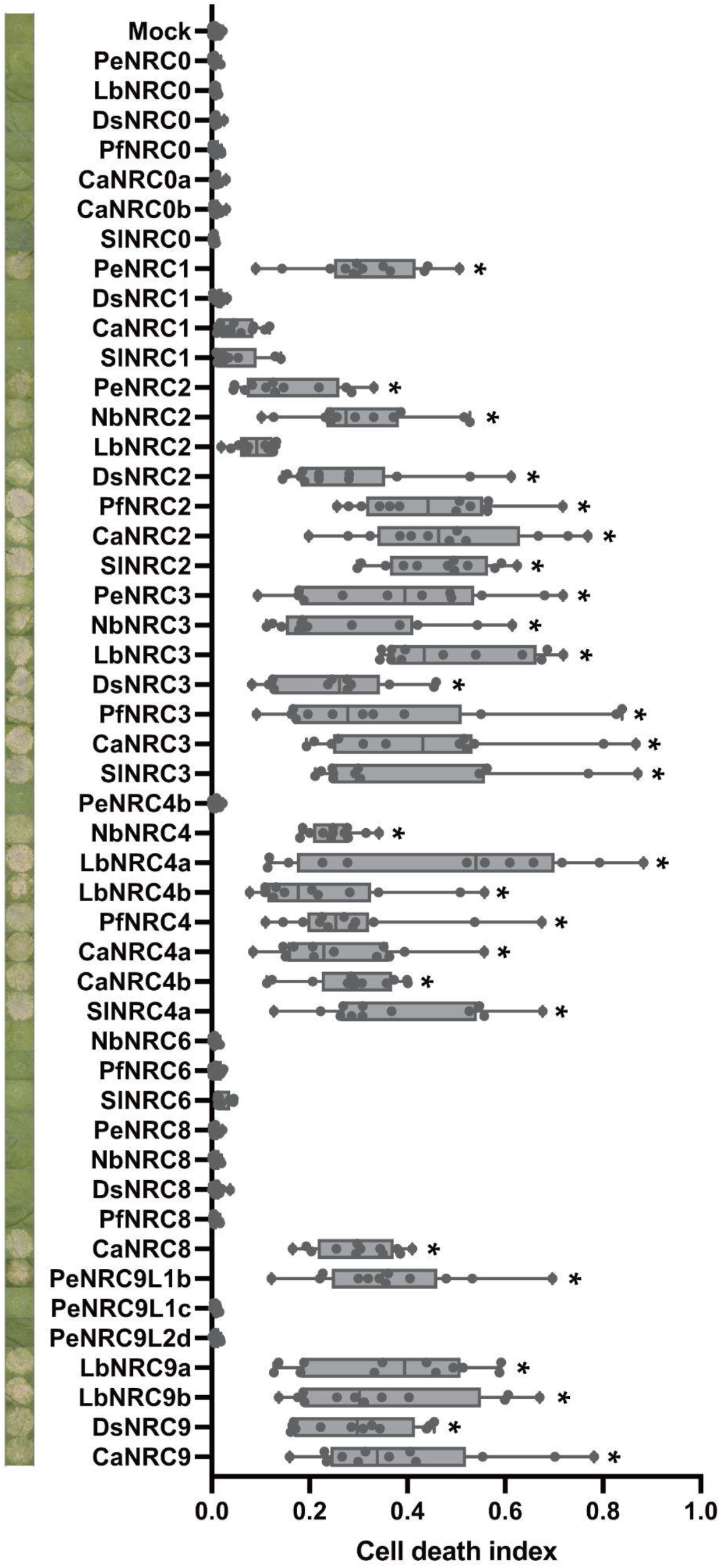
Cell death phenotypes and autofluorescence quantification following CP/Rx activation with solanaceous NRC helpers. Leaves of 27-day-old Nicotiana benthamiana nrc234 plants were agroinfiltrated with constructs expressing CP, Rx, and individual NRC helper variants. At 3 days post-agroinfiltration, infiltrated leaves were photographed and subjected to autofluorescence quantification. Multiple comparisons were performed using the Kruskal–Wallis test followed by Dunn’s post hoc test, with the P-value threshold set at 0.05. Groups labeled with asterisks indicate statistically significant differences.

**Supplemental Figure S4.**
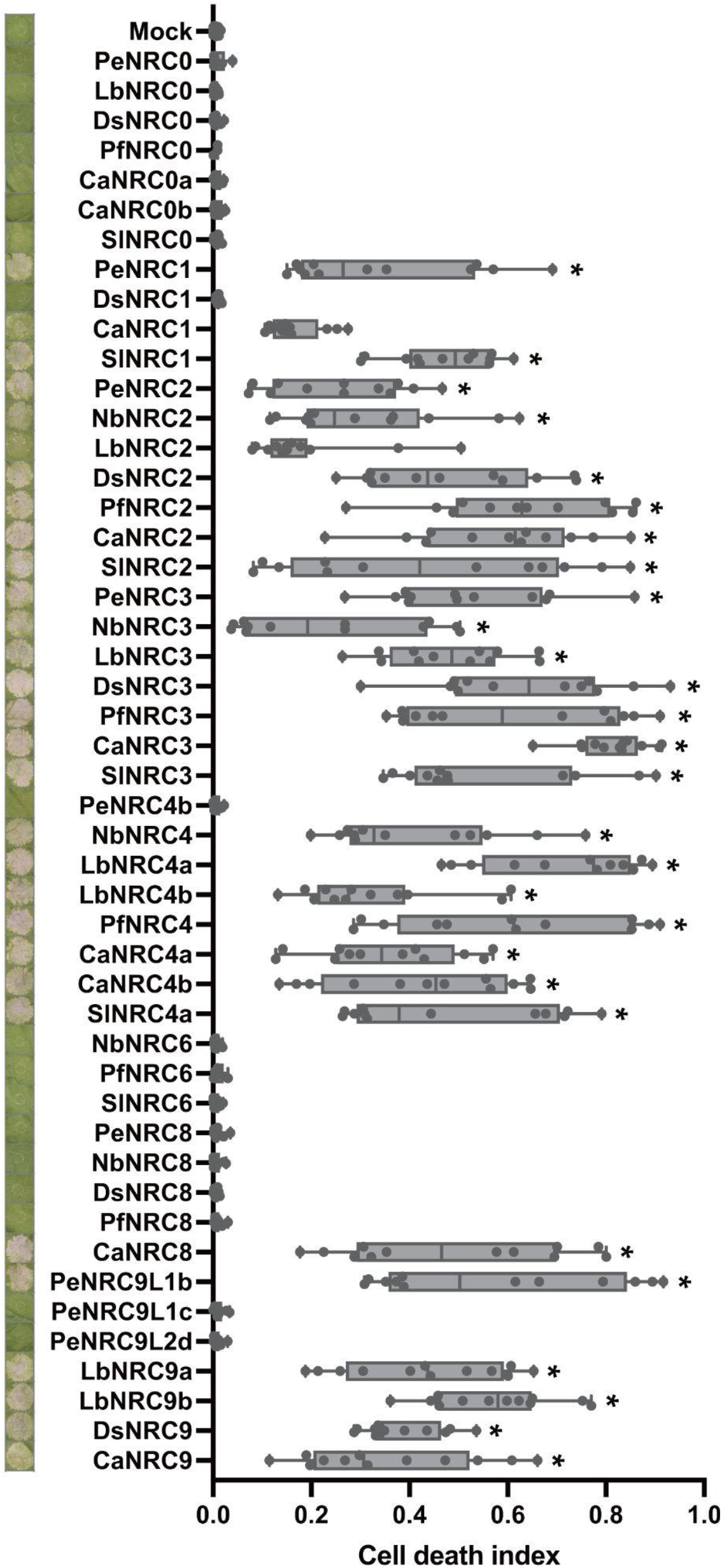
Cell death phenotypes and autofluorescence quantification following Avramr3/Rpi-amr3 activation with solanaceous NRC helpers. Leaves of 27-day-old *Nicotiana benthamiana nrc234* plants were agroinfiltrated with constructs expressing Avramr3, Rpi-amr3, and individual NRC helper variants. At 3 days post-agroinfiltration, infiltrated leaves were photographed and subjected to autofluorescence quantification. Multiple comparisons were performed using the Kruskal–Wallis test followed by Dunn’s post hoc test, with the P-value threshold set at 0.05. Groups labeled with asterisks indicate statistically significant differences.

**Supplemental Figure S5.**
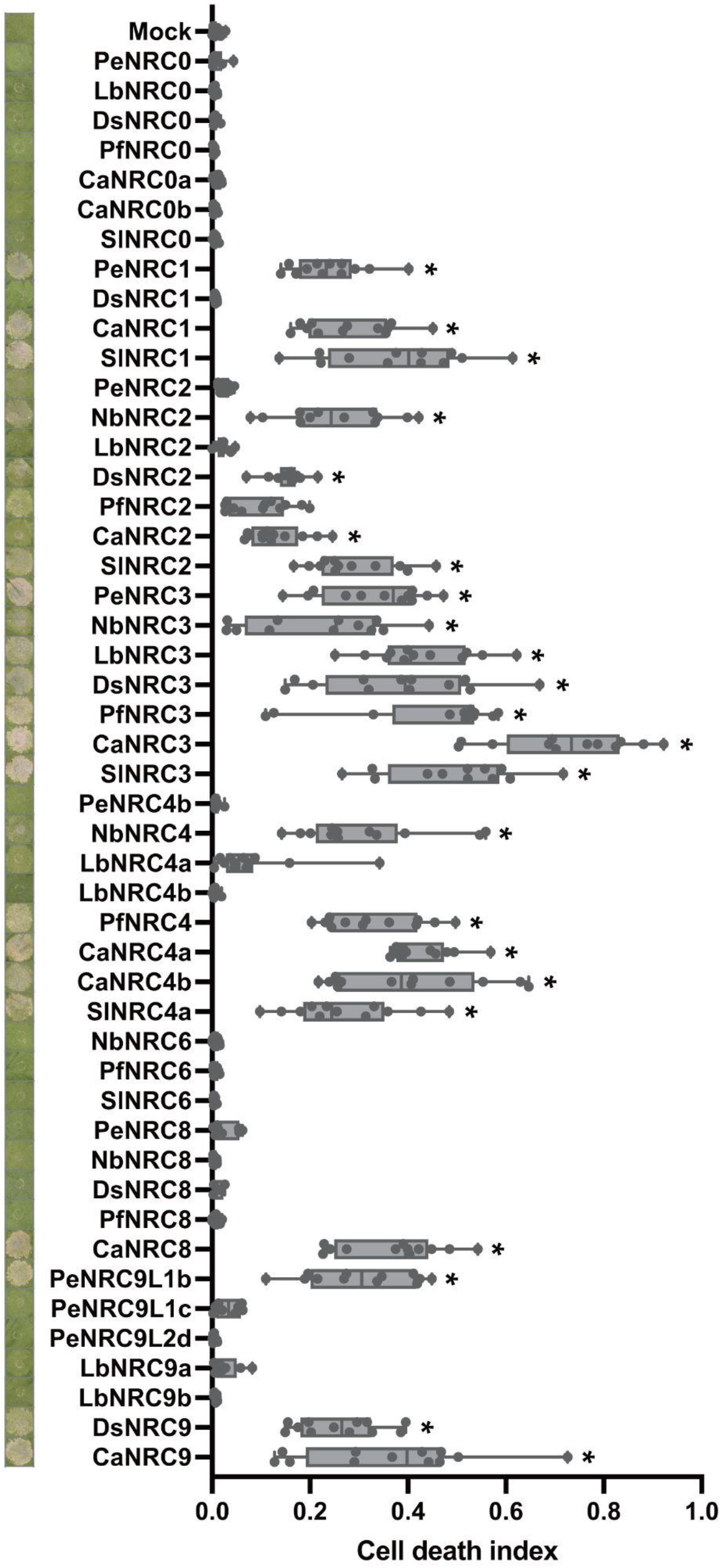
Cell death phenotypes and autofluorescence quantification following AvrBs2/Bs2 activation with solanaceous NRC helpers. Leaves of 27-day-old *Nicotiana benthamiana nrc234* plants were agroinfiltrated with constructs expressing AvrBs2, Bs2, and individual NRC helper variants. At 3 days post-agroinfiltration, infiltrated leaves were photographed and subjected to autofluorescence quantification. Multiple comparisons were performed using the Kruskal–Wallis test followed by Dunn’s post hoc test, with the P-value threshold set at 0.05. Groups labeled with asterisks indicate statistically significant differences.

**Supplemental Figure S6.**
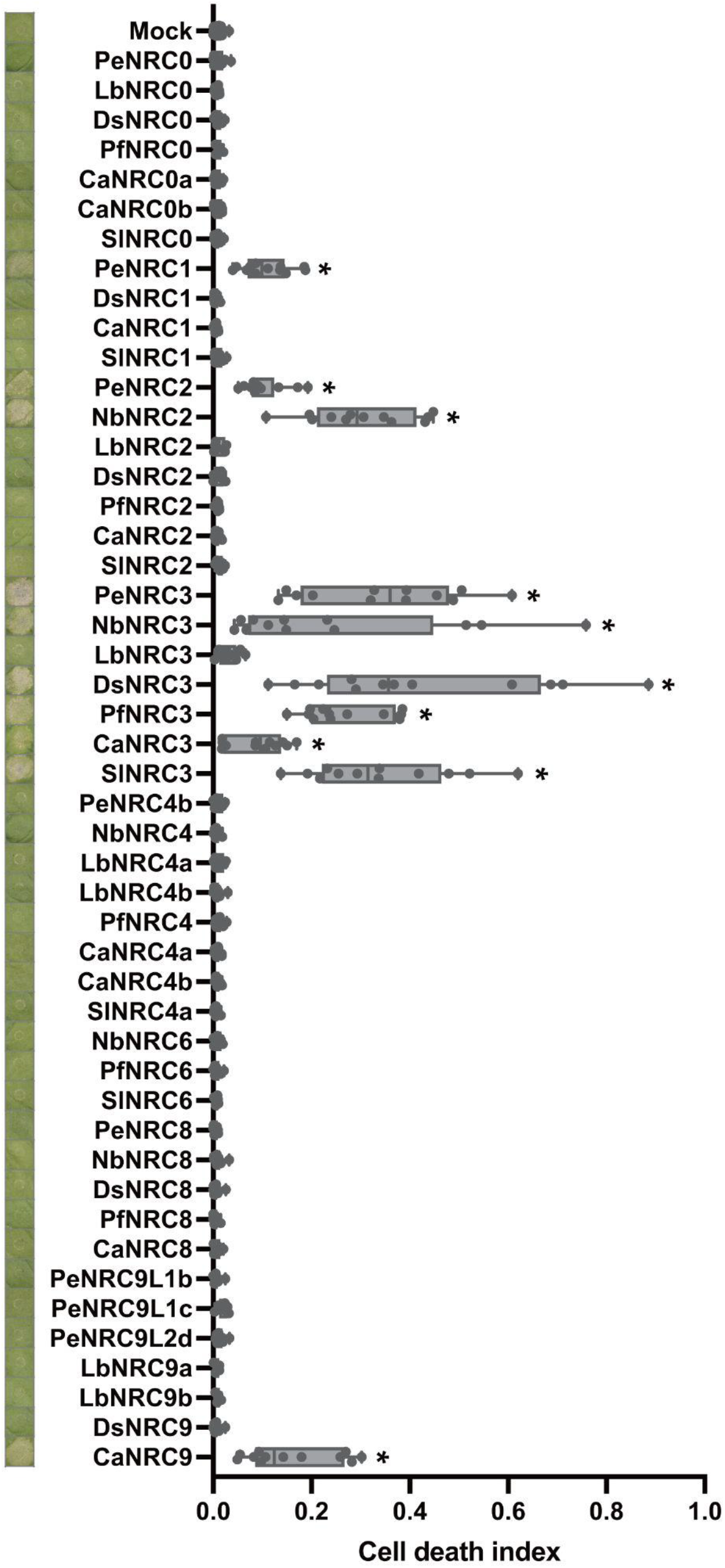
Cell death phenotypes and autofluorescence quantification following AvrPto/Prf activation with solanaceous NRC helpers. Leaves of 27-day-old *Nicotiana benthamiana nrc234* were agroinfiltrated with constructs expressing AvrPto, Pto, and individual NRC helper variants. At 3 days post-agroinfiltration, infiltrated leaves were photographed and subjected to autofluorescence quantification. Multiple comparisons were performed using the Kruskal–Wallis test followed by Dunn’s post hoc test, with the P-value threshold set at 0.05. Groups labeled with asterisks indicate statistically significant differences.

**Supplemental Figure S7.**
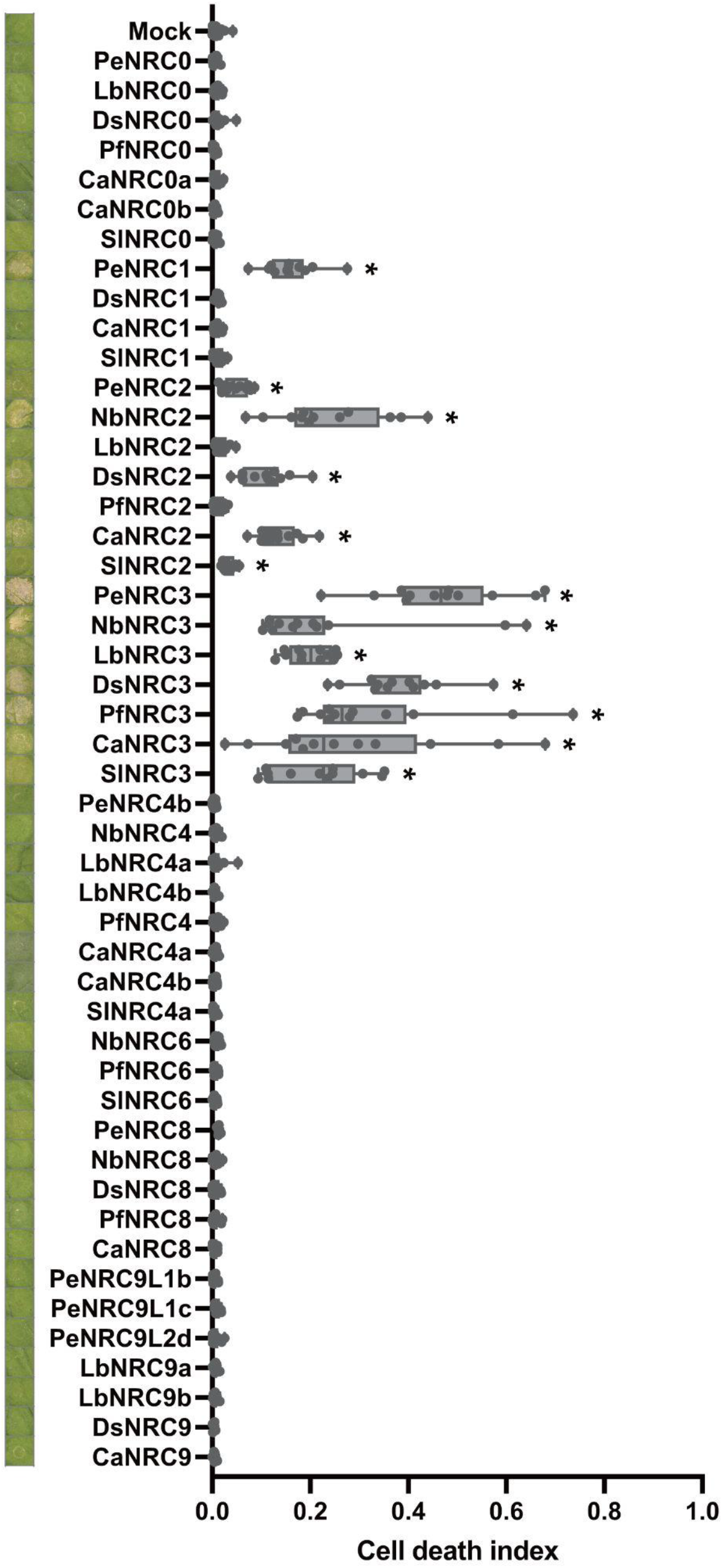
Cell death phenotypes and autofluorescence quantification following Avramr1/Rpi-amr1 activation with solanaceous NRC helpers. Leaves of 27-day-old *Nicotiana benthamiana nrc234* were agroinfiltrated with constructs expressing Avramr1, Rpi-amr1, and individual NRC helper variants. At 3 days post-agroinfiltration, infiltrated leaves were photographed and subjected to autofluorescence quantification. Multiple comparisons were performed using the Kruskal–Wallis test followed by Dunn’s post hoc test, with the P-value threshold set at 0.05. Groups labeled with asterisks indicate statistically significant differences.

**Supplemental Figure S8.**
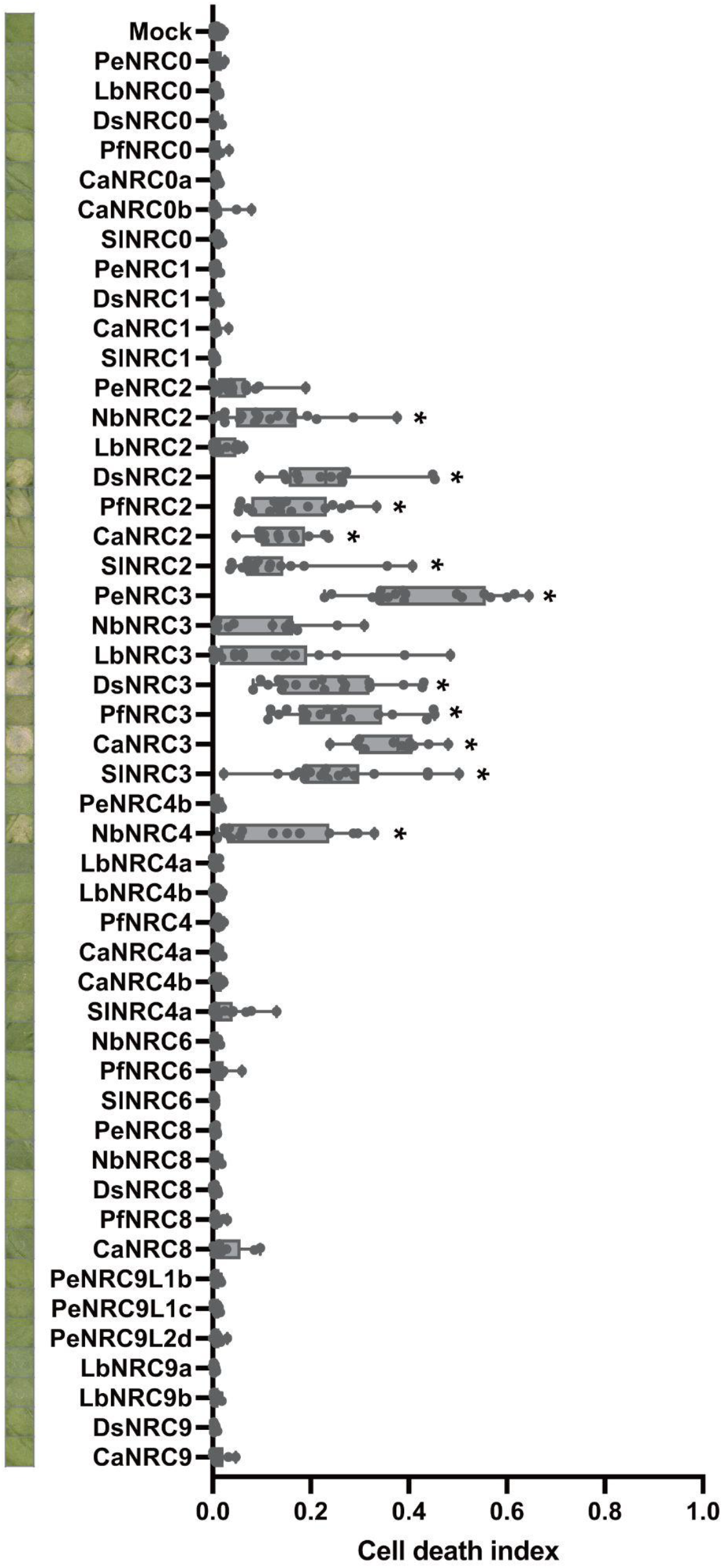
Cell death phenotypes and autofluorescence quantification following Nsm/Sw5b activation with solanaceous NRC helpers. Leaves of 27-day-old *Nicotiana benthamiana nrc234* plants were agroinfiltrated with constructs expressing Nsm, Sw5b, and individual NRC helper variants. At 3 days post-agroinfiltration, infiltrated leaves were photographed and subjected to autofluorescence quantification. Multiple comparisons were performed using the Kruskal– Wallis test followed by Dunn’s post hoc test, with the P-value threshold set at 0.05. Groups labeled with asterisks indicate statistically significant differences.

**Supplemental Figure S9.**
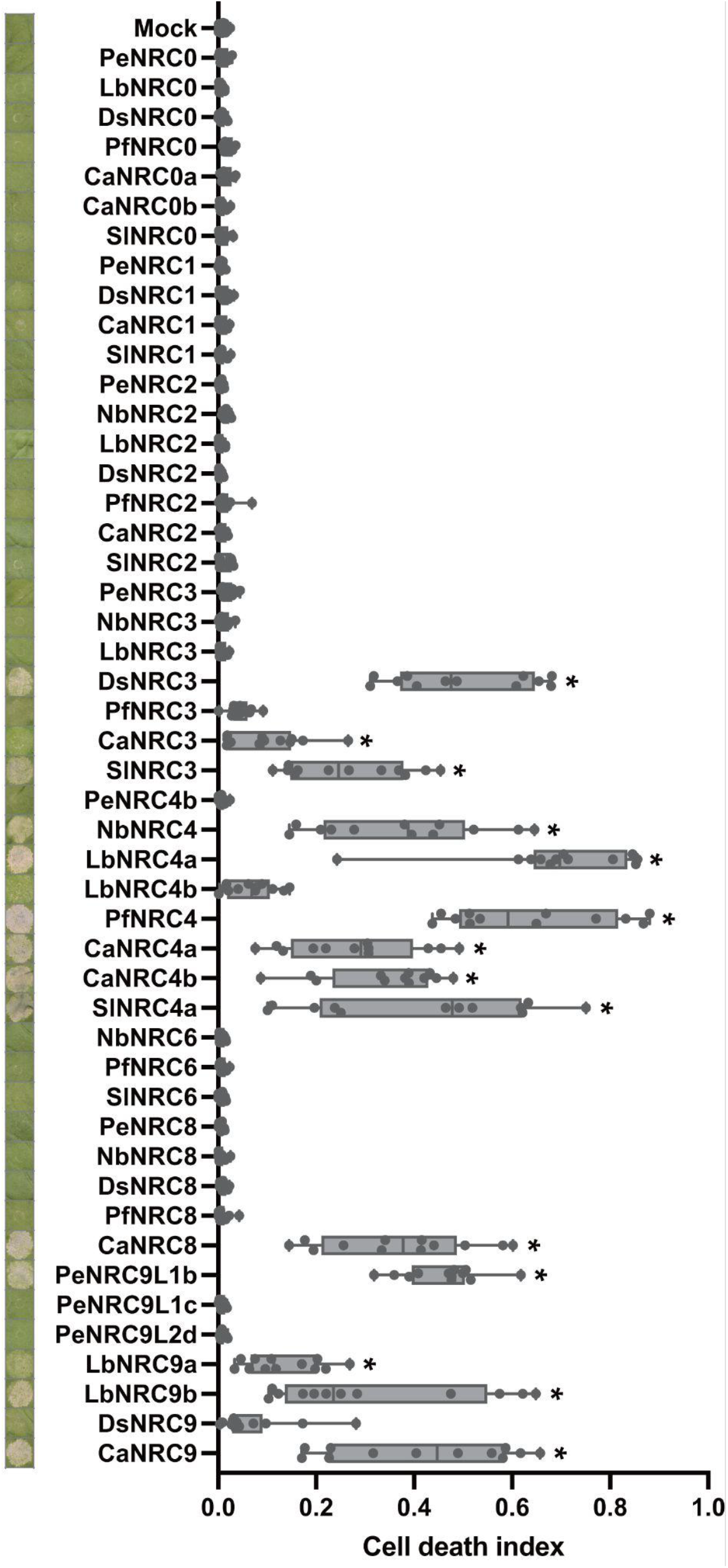
Cell death phenotypes and autofluorescence quantification following Avrblb2/Rpi-blb2 activation with solanaceous NRC helpers. Leaves of 27-day-old *Nicotiana benthamiana nrc234* were agroinfiltrated with constructs expressing Avrblb2, Rpi-blb2, and individual NRC helper variants. At 3 days post-agroinfiltration, infiltrated leaves were photographed and subjected to autofluorescence quantification. Multiple comparisons were performed using the Kruskal–Wallis test followed by Dunn’s post hoc test, with the P-value threshold set at 0.05. Groups labeled with asterisks indicate statistically significant differences.

**Supplemental Figure S10.**
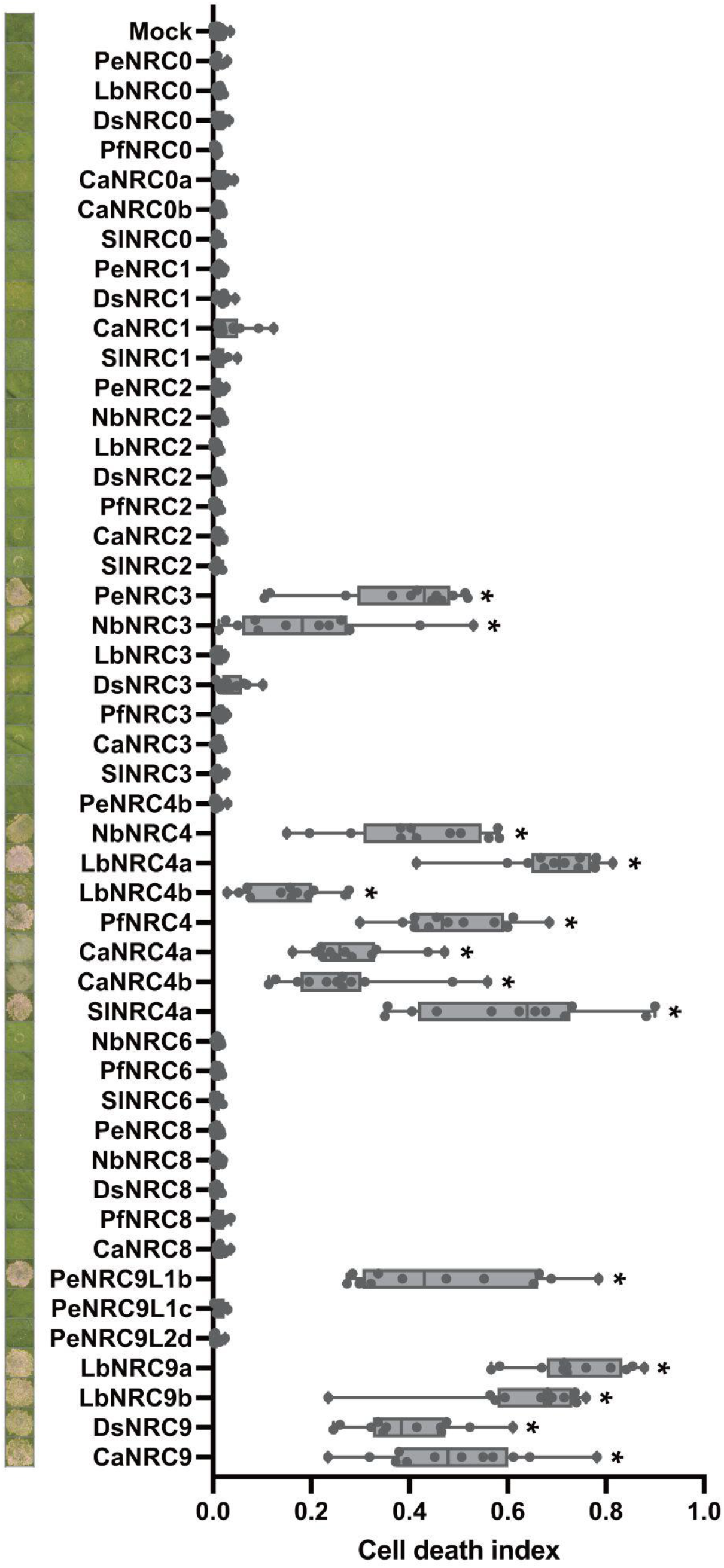
Cell death phenotypes and autofluorescence quantification following AVR1/R1 activation with solanaceous NRC helpers. Leaves of 27-day-old *Nicotiana benthamiana nrc234* were agroinfiltrated with constructs expressing AVR1, R1, and individual NRC helper variants. At 3 days post-agroinfiltration, infiltrated leaves were photographed and subjected to autofluorescence quantification. Multiple comparisons were performed using the Kruskal–Wallis test followed by Dunn’s post hoc test, with the P-value threshold set at 0.05. Groups labeled with asterisks indicate statistically significant differences.

**Supplemental Figure S11.**
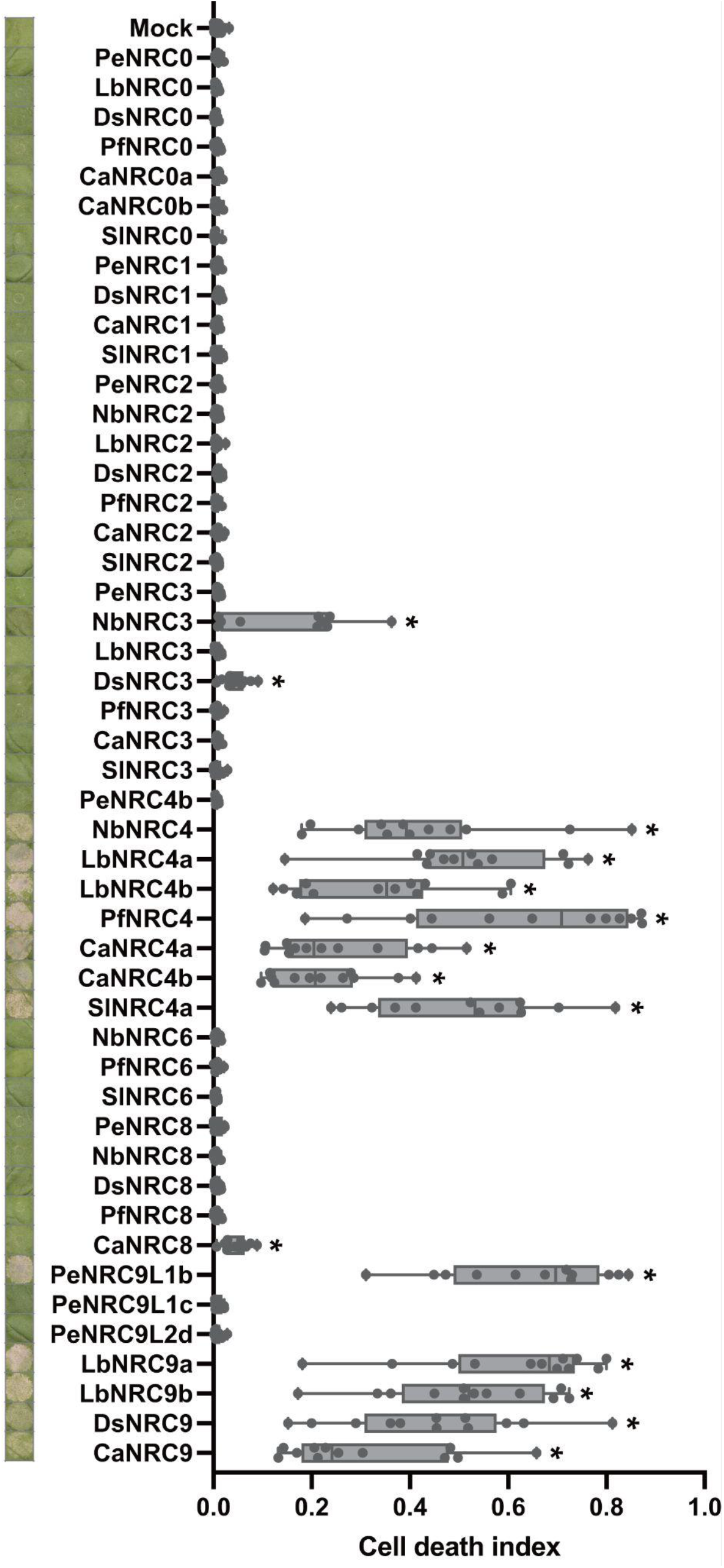
Cell death phenotypes and autofluorescence quantification following 9S activation with solanaceous NRC helpers. Leaves of 27-day-old *Nicotiana benthamiana nrc234* were agroinfiltrated with constructs expressing autoactive petunia 9S and individual NRC helper variants. At 3 days post-agroinfiltration, infiltrated leaves were photographed and subjected to autofluorescence quantification. Multiple comparisons were performed using the Kruskal–Wallis test followed by Dunn’s post hoc test, with the P-value threshold set at 0.05. Groups labeled with asterisks indicate statistically significant differences.

**Supplemental Figure S12.**
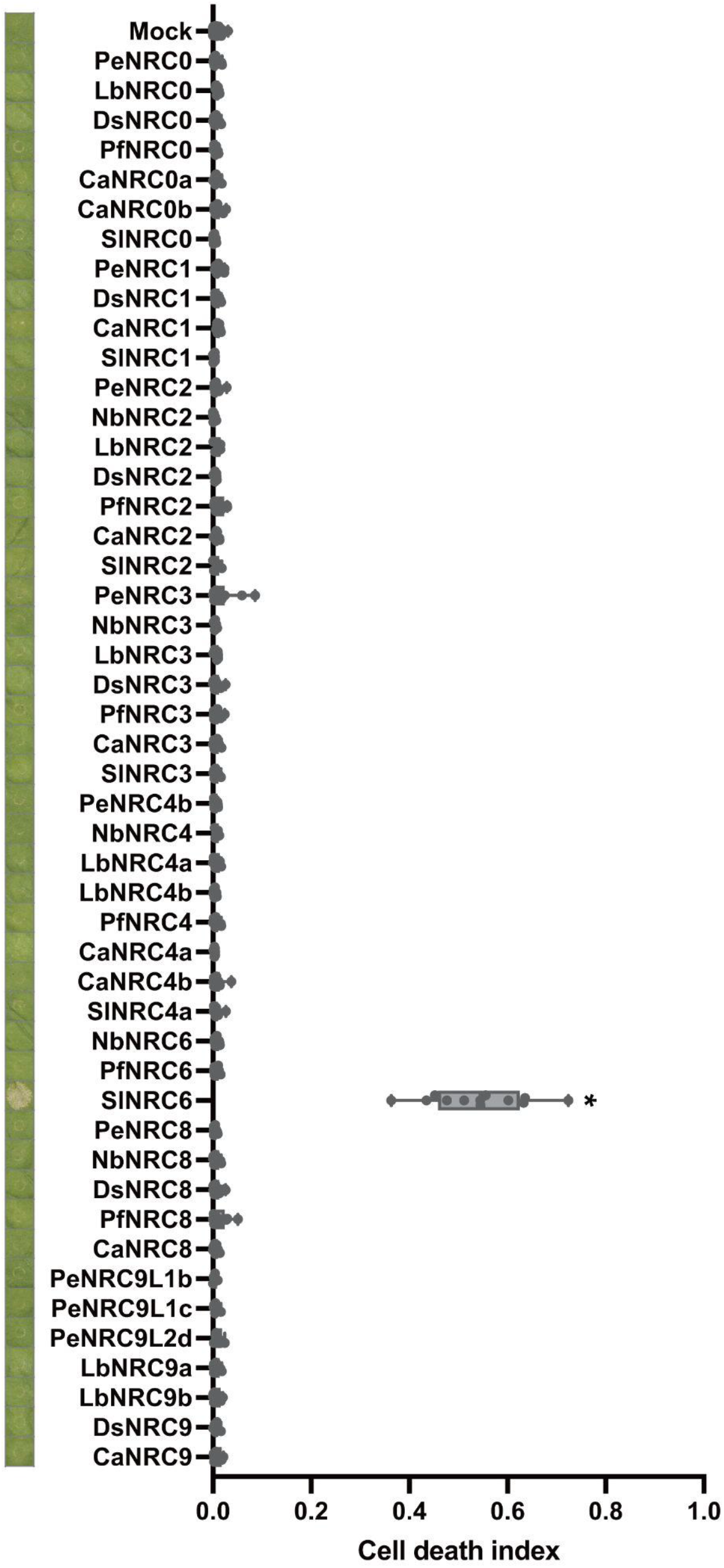
Cell death phenotypes and autofluorescence quantification following HeroF activation with solanaceous NRC helpers. Leaves of 27-day-old *Nicotiana benthamiana nrc234* were agroinfiltrated with constructs expressing autoactive HeroF and individual NRC helper variants. At 3 days post-agroinfiltration, infiltrated leaves were photographed and subjected to autofluorescence quantification. Multiple comparisons were performed using the Kruskal–Wallis test followed by Dunn’s post hoc test, with the P-value threshold set at 0.05. Groups labeled with asterisks indicate statistically significant differences.

**Supplemental Figure S13.**
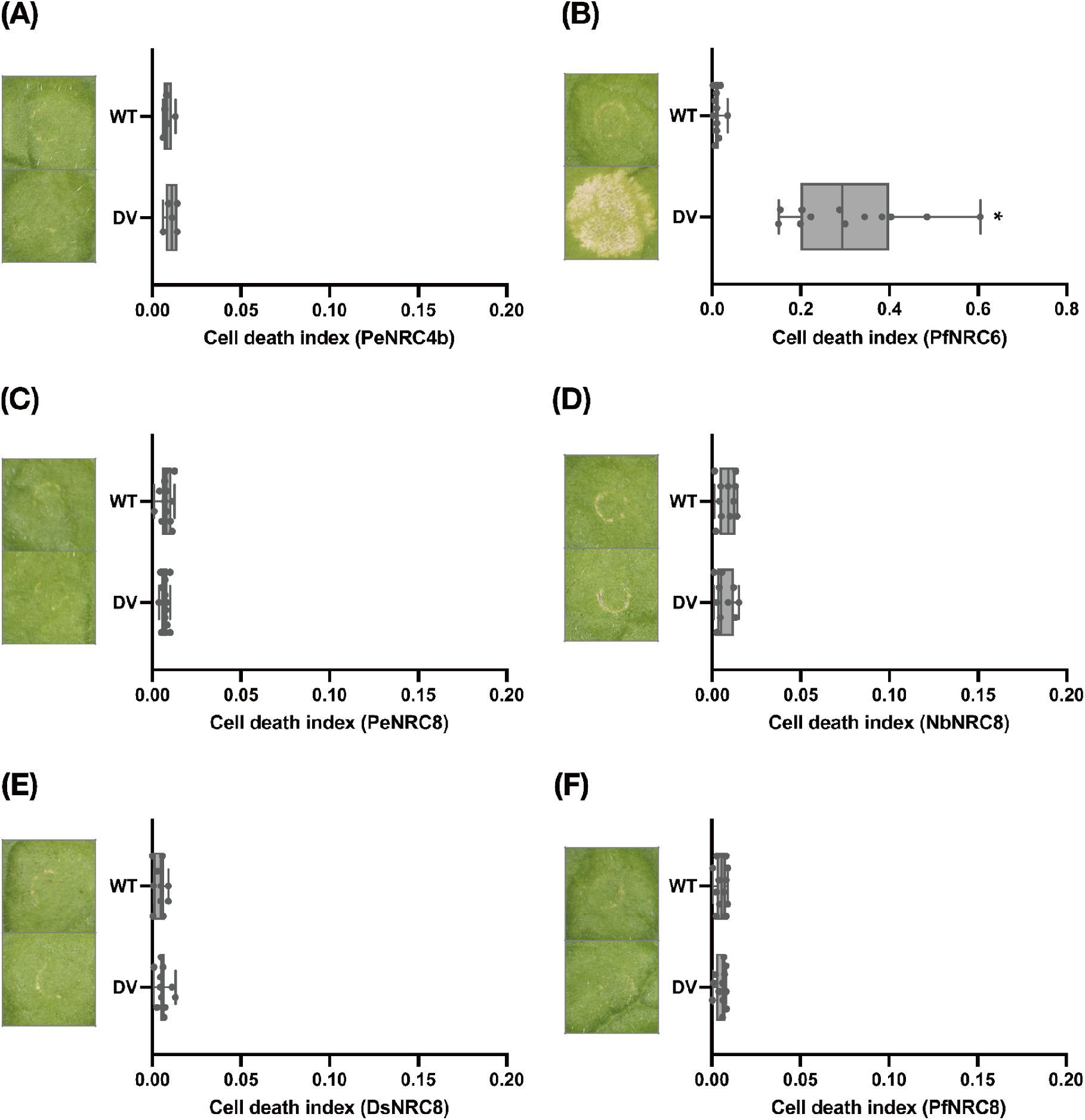
Comparison of cell death phenotypes and autofluorescence levels between wild-type and autoactive solanaceous NRC helpers. Leaves of 27-day-old *Nicotiana benthamiana nrc234* plants were agroinfiltrated with constructs expressing either wild-type or autoactive NRC helper variants. At 3 days post-agroinfiltration (dpi), infiltrated leaves were photographed and subjected to autofluorescence quantification. Multiple comparisons were performed using the Kruskal–Wallis test followed by Dunn’s post hoc test, with the P-value threshold set at 0.05. Groups labeled with asterisks indicate statistically significant differences. (A) PeNRC4b; (B) PfNRC6; (C) PeNRC8; (D) NbNRC8; (E) DsNRC8; (F) PfNRC8.

**Supplemental Figure S14.**
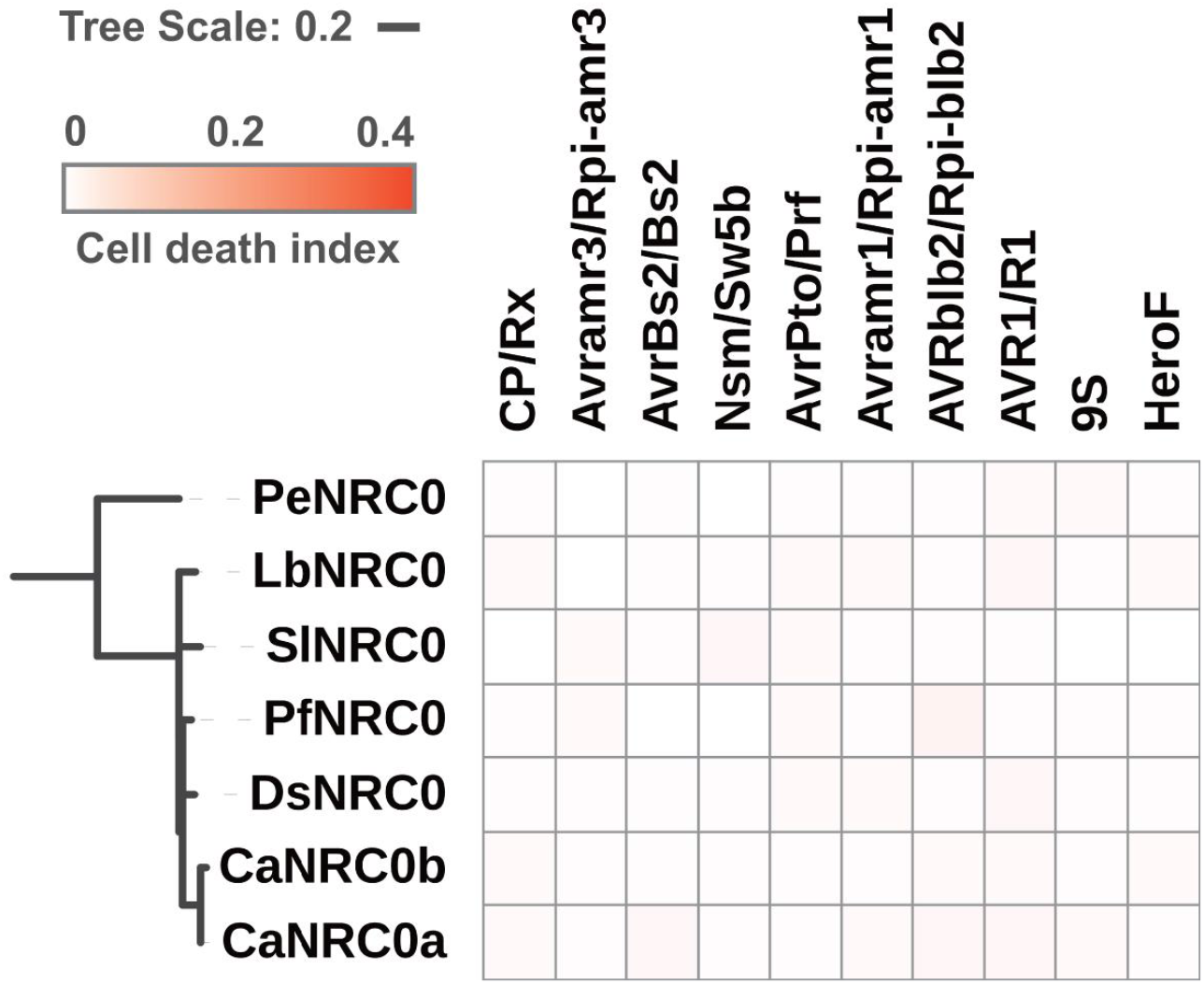
Functional compatibility matrix of solanaceous NRC0 with diverse effector–sensor NLR pairs. Cell death assays were conducted in *nrc234* knock out *N. benthamiana* mutants triggered by pairing each NRC helper variant from seven Solanaceae species (petunia, *N. benthamiana*, goji berry, jimsonweed, lantern fruit, pepper, and tomato) with a panel of canonical effector–sensor NLR combinations. Tested pairs include CP/Rx, Avramr3/Rpi-amr3, AvrBs2/Bs2, Nsm/Sw5b, AvrPto/Prf, Avramr1/Rpi-amr1, AVRblb2/Rpi-blb2, AVR1/R1, and two autoactive sensors (petunia 9S and tomato HeroF). Heatmaps summarize the degree of cell death responses across NRC0–sensor combinations. Robust cell death, weak cell death, and no response are indicated by graded color intensities. The phylogeny trees of NRC helpers were subsetted from the master tree of NRC helpers (Fig. 1D).

**Supplemental Figure S15.**
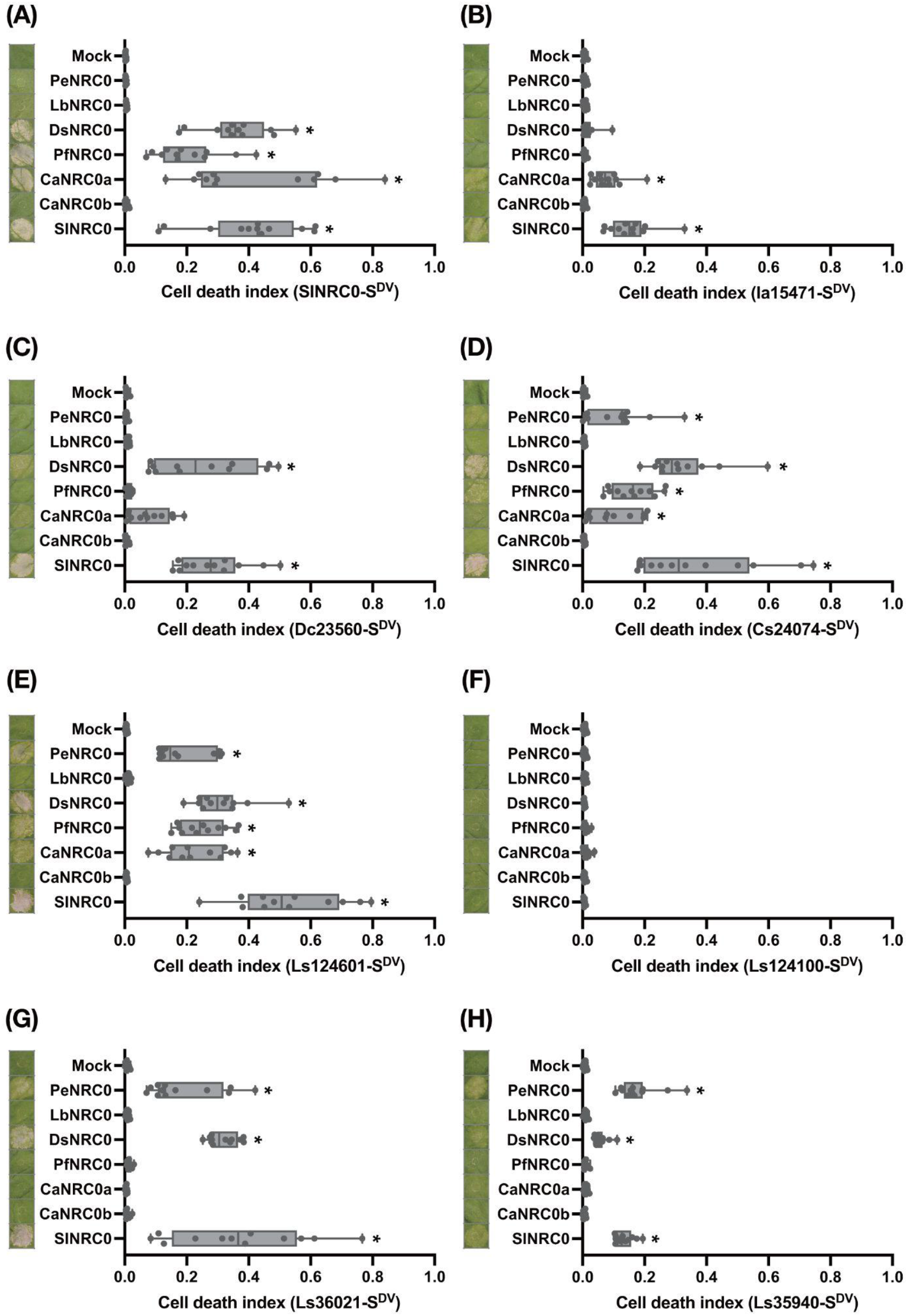
Cell death phenotypes and autofluorescence quantification following the activation of NRC0-dependent sensors with NRC0 variants. Leaves of 27-day-old *Nicotiana benthamiana nrc234* were agroinfiltrated with constructs expressing autoactive NRC0-dependent sensors and individual NRC0 variants. At 3 days post-agroinfiltration, infiltrated leaves were photographed and subjected to autofluorescence quantification. Multiple comparisons were performed using the Kruskal–Wallis test followed by Dunn’s post hoc test, with the P-value threshold set at 0.05. Groups labeled with asterisks indicate statistically significant differences. (A) SlNRC0-S^DV^; (B) Ia15471-S^DV^; (C) Dc23560-S^DV^; (D) Cs24074-S^DV^; (E) Ls124061-S^DV^; (F) Ls124100-S^DV^; (G) Ls36021-S^DV^; (H) Ls35940-S^DV^.

